# Microtubule-Inspired Functionalization of Carbon Nanotubes: A Biomimetic Carrier Design

**DOI:** 10.1101/2022.01.20.477082

**Authors:** Karina de Almeida Barcelos, Laleh Alisaraie

**Affiliations:** School of Pharmacy, Memorial University of Newfoundland, 300 Prince Philip Drive, A1B 3V6, St. John’s, Newfoundland, Canada

**Keywords:** Carbon nanotube, Functionalization, Bioavailability, Microtubule, Lateral and longitudinal sites, Antimitotic agents, Computational simulations

## Abstract

We propose a bioinspired, non-covalent carbon nanotubes (CNTs) functionalization strategy to augment their bioavailability and alleviate their biotoxicity. For functionalization, select amphiphilic peptides from a cytoskeletal biopolymer, microtubule (MT), were used. The peptides are involved in the MT polymerization by maintaining the essential lateral interactions among the MT’s α- and β-tubulin subunits. They also participate in forming the MT-binding sites for hosting several MT-targeting antimitotics. Utilizing *in silico* methods, this study showed the peptides influenced CNT’s diffusivity and aqueous solubility. The hydrodynamic shield formed by the peptides from β-tubulin was more widespread on the CNT than the α-tubulin peptides’; however, the latter created a broader hydrophobic CNT coating than those from the β-tubulin. In particular, the peptides consisting of the H1-B2, H10, H1-B2, and the M-loop, demonstrated structural features that serve to augment CNTs’ water solubility and dispersibility. The performance of the peptide-functionalized CNTs as drug carriers was examined by studying seventeen antimitotics. The CNT-peptides structural composition was identified as a suitable carrier for phomopsin A, laulimalide, epothilone A, epothilone D, discodermolide, eribulin, and docetaxel. The peptides played dual roles displaying affinities to the antimitotics and the CNT; in particular, the peptides from the H1-B2 and H2-B3 loops of β-tubulin exhibited exceptional binding properties. Specific mutations on the wildtype peptides, including those from the α-tubulin M-loop and H2-B3, or the β-tubulin H1-B2, are proposed to refine their hydrophobicity, eliminate unfavorable inter-peptides electrostatic interactions or the spatial hindrance at certain regions, to enhance their conformational steadiness and exposure to the tube surface. A combination of the select amphiphilic peptides from both tubulin subunits is suggested to improve CNTs bioavailability and efficiency for carrying insoluble hydrophobic cargos.

## 1. Introduction

There have been emerging interests in exploring carbon nanotubes (CNTs) physicochemical, mechanical, thermal, structural, and optical properties^1^ for their applications in multiple fields, including nanomedicine^2^, drug, and gene delivery. ^3-12^ The pristine CNTs’ chemical structures comprise carbon atoms with sp^2^ hybridization, capable of interacting with other chemical entities through van der Waals (vdW) forces. CNTs accumulate in cancer tissues due to their enhanced permeability and retention (EPR) properties, which besides their high aspect ratio and unique architecture, make them potential nanoparticles for onsite cancer drug delivery systems.^13^ Their hydrophobic property allows their bound hydrophobic cargos (e.g. drugs) to float in the bloodstream. CNTs high aspect ratio enables a high payload that consequently would help reduce the required drug dose for cancer treatments ^14^ by lowering the potential off-target drug interactions and the associated adverse effects in cancer patients.^15, 16^ However, a significant obstacle for CNTs’ bio-applicability is their low aqueous solubility.

Some chemical functionalization methods have been developed to address the probelm^17^, including covalently attaching hydrophilic groups, usually carboxylic acid or amide chemical functional groups, via oxidation routes or using any existing defects in the CNT structure, or introducing artificial holes for chemical substitutions.^18^ Carboxylation of CNTs has been shown to improve their oxidative degradation process, potentially reducing their toxicity. The particles produced from the oxidative degradation could be more easily expelled from the respiratory system than the unprocessed nanotubes.^19^ Similarly, oxidation of multi-walled carbon nanotubes (MWCNTs) using H_2_SO_4_ and HNO_3_ results in O-MWCNTs, which have been shown to not only improve CNTs biodegradation *in vitro* but also could act as an anticancer cytotoxic agent.^20^ While surface oxidation of CNTs could increase their biodegradation *in vitro* and *in vivo*, assisting in the excretion of the nanoparticles, it could also interrupt CNT’s function as a drug delivery system when they immaturely release their bound cargos (i.e. drugs) before delivery to the site of action.^21^ Despite the effectiveness of the covalent methods on CNTs hydrophilicity, they disrupt the intrinsic sp^2^ hybridization of many carbon atoms and their associated network that are necessary for the CNT’s exceptional properties^1^ and thus diminish their utility for particular applications.^17^ Alternatively, physical surface-adsorption has gained researchers’ interest for maintaining the sp^2^ hybridization in the chemical skeleton of carbon-based nanomaterials such as CNT^22, 23^, graphene^22, 24^, and fullerene.^22, 25^ Amphiphilic peptides consisting of amino acids with various degrees of side-chain hydrophobicity and hydrophilicity are suitable molecules that due to their dual binding properties could bind to CNTs and water molecules.

The presented work proposes a bioinspired strategy for improving bioavailability and biocompatibility of single-walled carbon nanotubes using some of the functionally important amino acids sequence segments of the cytoskeletal biopolymer, microtubule (MT), for CNTs non-covalent functionalization. MTs are composed of α-β tubulin heterodimer building blocks. MTs regulate multiple vital cellular functions and act as a platform for motor proteins (e.g., dynein^26, 27^, and kinesin^28^) movements for intracellular cargo transportations. The MTs’ continual dynamic reconstruction is a natural process that permits unceasing transitions between heterodimers assembly (polymerization and elongation) and disassembly (depolymerization and shrinkage) at the MT ends^29^. The extension of MTs results from the heterodimers longitudinal (head-to-tail) assembly via non-covalent interactions of the α- and β-tubulin subunits that form long chains of protofilaments (pfs). The subunits of the pfs interact laterally in side-by-side arrangements, shaping a hollow cylinder along MT’s longitudinal axis^30-35^.

Due to the various and vital roles of the MTs in cell formation, cargo and organelle transportations, and cell motility, they have been one of the important targets for drug discovery and development of antimitotic, anticancer drugs.^36^ The drugs act either as an MT polymerizer or depolymerizer, inhibiting its dynamic assembly-disassembly (i.e., polymerization-depolymerization) process, essential for maintaining the cell structure and survival. There are several binding sites on the α-, β-tubulin, located at the lateral or longitudinal interface of the two tubulin monomers that could accommodate several antimitotics, such as paclitaxel^37^, docetaxel^38^, taccalonolide^39^, epothilones (types A, B, D, and ixabepilone)^40, 41^, cyclostreptin^42^, dictyostatin^43^, discodermolide^44^, zampanolide^45^, laulimalide^46, 47^, peloruside A^46, 47^, vinblastine^48-50^, dolastatin^51^, phomopsin A^52^, and eribulin^53^.

Earlier *in vivo* studies have suggested that MWCNTs mimic some of the MT’s properties in HeLa cells. It reduces MT’s dynamic, affecting the protein’s polymerization-depolymerization process and its performance during cell division^54^. Due to the high similarity of CNTs and MTs^55^ and their potential for use as drug delivery system ^56^, this work aims to take a step towards biomimicry of the MT using CNT to project some of the MT’s drug-binding properties for improving CNTs drug delivery properties. In this approach, the MT segments from α-β tubulin heterodimer were selected as the amphiphilic peptides capable of non-covalently binding to the CNT surface. The peptides participate in the functionally critical lateral and longitudinal interactions among tubulin subunits, the MT protofilaments formation, and its dynamic “treadmilling” processes^37^. The peptides also contribute to the construction of the MTs drug binding sites.

The proposed microtubule-inspired CNT functionalization in this study is expected to enhance its solubility and drug delivery potential. The chemical specificity and amphiphilic diversity of the selected MT-based peptides can weaken CNTs’ biotoxicity and amplify their cellular uptake.^57^ *In silico* methods were utilized for this investigation to acquire information on the molecular and atomic interactions between the peptides bound to the CNT surface and each of the seventeen aforementioned antimitotic MT-targeting drugs. This work examined the potential of the proposed biomimicry strategy for improving CNTs application as biocompatible nanocarriers.

## 2. Materials And Method

There are five main lateral-associated segments on each α- and β-subunit of tubulin in the crystal structure of the α-β tubulin heterodimer.^37^ They are located at: (*i*) H1-B2 loop (α: Tyr24–Pro63, β: Ile24–Ala65); (*ii*) H2-B3 loop (α: Arg79–Pro89, β: Gly81–Asn91); (*iii*) H4– T5 loop (α: Leu157–Tyr161, β: Glu159–Asp163); (*iv*) M-loop (α: Tyr272–Val288, β: Pro274– Glu290); and (*v*) H10 helix (α: Asp327–Ile341, β: Asp329–Phe343). The H10 helix is the only MT fragment that participates in the α- and β-subunits’ both lateral and longitudinal interactions. The amino acids indices and the protein segments’ secondary structure (SS) assignments are according to the crystal structure of α-β tubulin heterodimer of *Bos taurus* (Uniprot code, P81947 (α-tubulin) and Q6B856 (β-tubulin)), deposited on the Protein Data Bank^58^ under 1JFF.pdb^37^ retrieving code. Here, 1JFF.pdb^37^ (*Bos taurus*) was studied instead of *Homo sapien* tubulin subunits due to the availability of the experimentally-solved structure of the former. The sequence similarities between pair of equivalent peptides of α- and β-tubulin (i.e., the P1-P6; the P2-P7; the P3-P8; the P4-P9; and the P5-P10) were calculated using the Lalign server.^59, 60^ (**Table 1** and **Figure 1**)

**Table 1:**
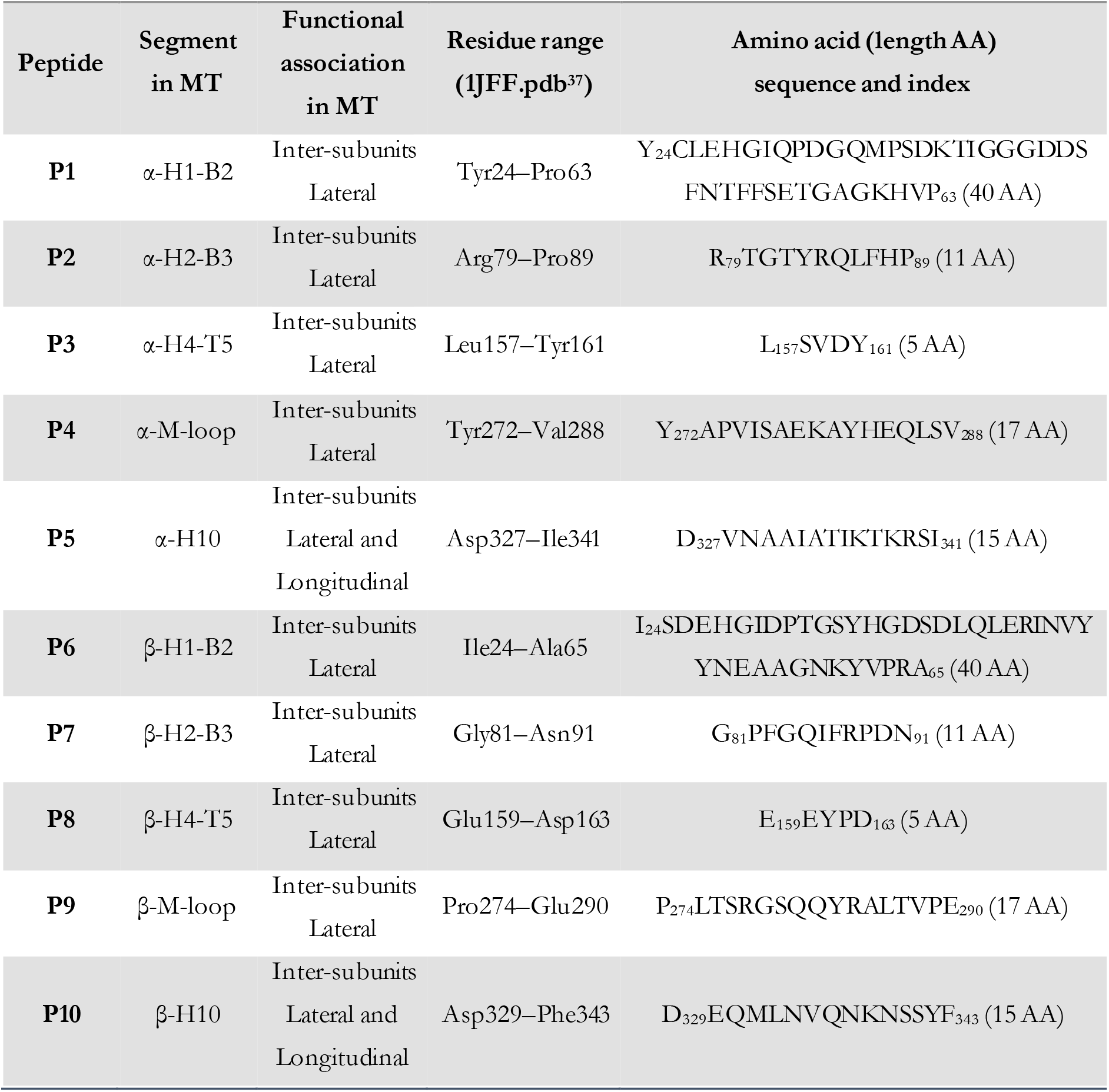
The studied MT segments (i.e., the P1–P10 peptides) with their assigned SS on the tubulin heterodimer crystal structure (1JFF^37^) and location in an MT protofilament. The amino acids are shown as single-letter codes. The letters “T”, “B”, and “H” refer to the loops, β-strand, and α-helix folding, respectively.

**Figure 1:**
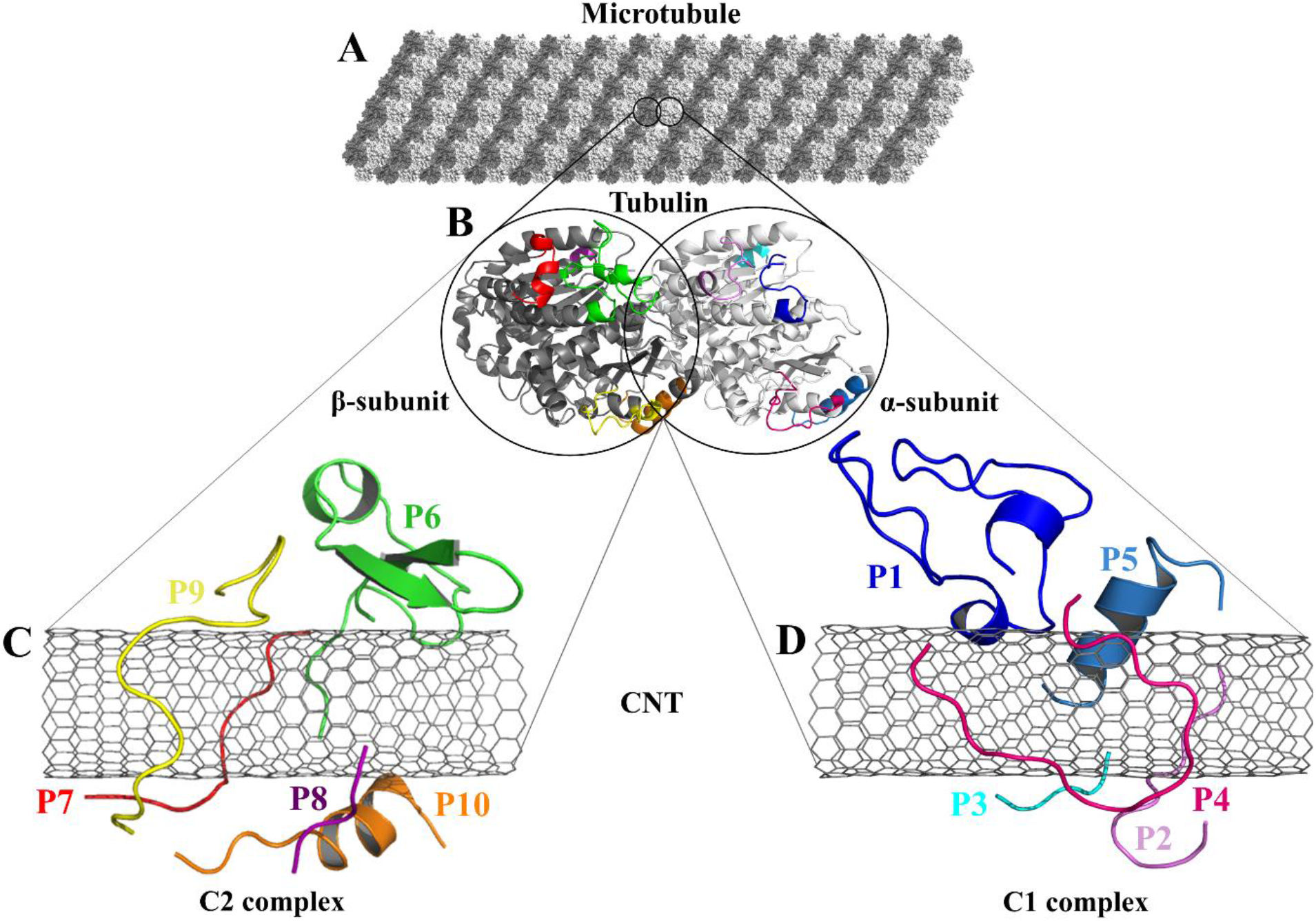
(**A**) The schematic of an MT composed of (**B**) the tubulin heterodimers with α-subunit (light grey), β-subunit (dark grey) (1JFF.pdb^37^), and the studied tubulin-segments as loaded peptides on the CNT, (**C**) the C2 system consists of the peptides from β-tubulin segments and (D) the C1 system of the peptides from α-tubulin.

The most critical lateral interactions among the MT protofilaments are directed by conformational changes of the tubulin fragments that consist of the M-loop (the P4 and P9) and the H1-B2 loop (the P1 and P6).^37, 61^ The highly flexible loops undergo a wide range of changes at their dihedral angles leading the MT lattice to shape MTs with different diameters by the association of 12–16 interacting protofilaments. ^30, 34^ The M-loop plays a critical role in the pfs formation and accommodating anticancer drugs. ^34, 37, 61-63^ (**Table 1**)

### 2.1. In Silico Preparation of the Peptides

The missing (i.e. unsolved) amino acids from the crystal structure of α-β tubulin heterodimer 1JFF.pdb^37^ were identified according to the tubulin’s amino acid sequence information. They were modeled and inserted in the heterodimer structure^35^. The protein fragments were next extracted and named the peptides P1–P5 (from α-tubulin) and P6–P10 (from β-tubulin). (**Table 1**)

#### 2.1.1. In Silico Preparation of the C1 and C2 Structures

CNTs are specified by their diameter and chiral angle, determined with (n,m) vector pair notation. The initial three-dimensional structure of the single-walled CNT was modeled using Nanotube Modeler (version 1.8.0, JCrystalSof)^64^ in a zigzag index of (16,0) with 12.4 Å in diameter and 40.0 Å in length, comparable to our previous experiments.^65^ The zigzag CNT (16,0) also has sufficient surface area for loading all the selected peptides, the P1–P10, onto the CNT’s outer wall. (**Table 2**)

**Table 2:** Description of the CNT-peptides complex systems.

| Peptides-CNT Complex System | System descriptions | Peptides | Chirality | No. of CNT |
| --- | --- | --- | --- | --- |
| **C1** | P1–P5 and | n = 5 | Zigzag | n = 1 |
| | CNT (16,0) | from α-tubulin (the P1 to P5) | (16,0) | |
| **C2** | P6–P10 and | n = 5 | Zigzag | n = 1 |
| | CNT (16,0) | from β-tubulin (the P6 to P10) | (16,0) | |

The P1–P5 (in the C1 system) and the P6–P10 (in the C2 system) were positioned randomly over the length of the CNT surface at a distance > ∼3 Å. Next, the initial structures of the C1 and C2 were subject to molecular dynamics (MD) simulations as follows:

### 2.2. Parametrization and Set-up for the MD Simulations

The MD simulations were carried out using the Gromacs package v.2016.5 ^66, 67^. The OPLS-AA force field^68^ was employed for the calculation of bonding and non-bonding interactions.^65^ The CNT non-bonded and charging parameters were according to the configurations described by Li *et al*.^65^, whereas the peptides’ topology parameters were obtained from the Gromacs topology library for amino acids. (**Table 3**)

**Table 3:**
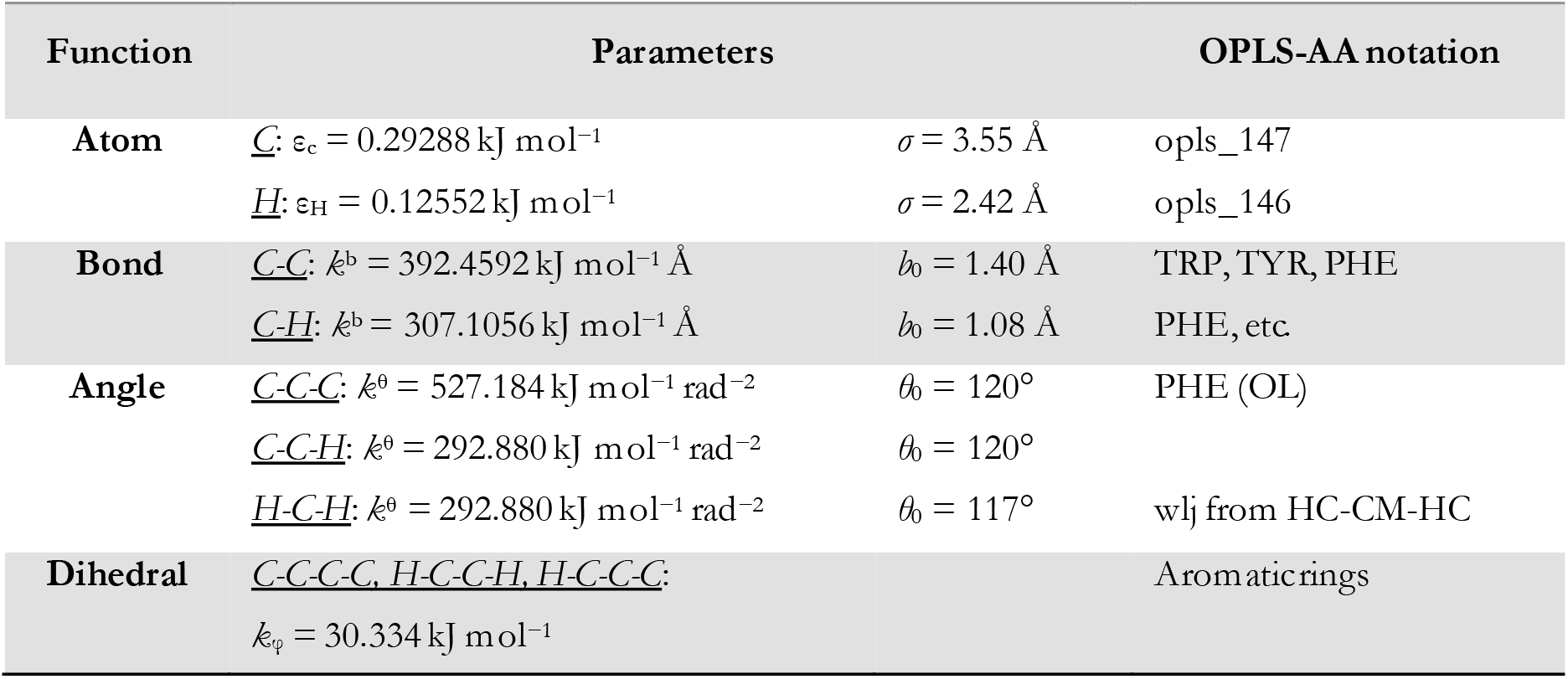
Non-bonded parameters for the MD simulation of pristine CNT according to the OPLS-AA force field. The *k*^*b*^, *k*^*θ*^, and *k*_*φ*_ are the force constants of stretching, bending, and torsional potentials, respectively; *b*_*0*_ and *θ*_*0*_ are reference geometry parameters; *ε* is the energy well-depth, and *σ* is the separation distance at which the interparticle potential is zero. Partial charges for carbon and hydrogen atoms were set 0.0, as for the uncharged particles.^65^

Besides the C1 and C2 systems, a separate MD simulation was performed (as a control system) for a single pristine CNT (16,0), in the absence of the peptides in a cubic water box with the same dimensions as in the C1 and C2 simulations (XYZ dimensions: 8.0 × 8.0 × 8.0 nm^3^). In addition, ten other independent MD simulations were carried out in the absence of the CNT for each of the ten peptides (i.e., a free, unbounded peptide in a water box), which were also considered control systems. The latter is labeled with a single-prime symbol (‘) as in the P1’ to P10’ to distinguish each CNT-bound from the free peptide. (**Table S1**)

The simple point charge (SPC) water model was used for modeling the solvent (water) box. Depending on the resulting net charge of each system, Na^+^ or Cl^−^ were added as counter ions to attain a 0.0 net charge under the physiological pH. (**Table S2**)

Periodic boundary conditions (PBC) were applied in three dimensions to minimize the edge effects by replicating adjacent boxes in all directions. Gromacs used the minimum image convention to ensure that only one image of an atom is examined for a short-range pair interaction during the calculations under the PBC. The non-bonded vdW and electrostatic interactions were modeled according to the forcefield parameters and 1.4 nm cut-off distance for both interaction types. The electrostatic forces and energy calculations were based on the Particle Mesh Ewald (PME) algorithm^69^ to obtain accurate long-range interactions. Energy minimization (EM) was performed on the C1, C2 systems, the free peptides, and the pristine CNT using the steepest descent (SD)^70^ algorithm with a maximum force tolerance ranging from 100.0–1000.0 kJ mol^−1^ nm^−1^. (**Table S3**)

The energy minimization was followed by the position restraint (PR) using the “md” integrator for implementing Newton’s equation of motion for 4.0 ns duration with 2.0 × 10^−3^ ps time-step. The Linear Constraint Solver (LINCS)^71^ algorithm was applied to constrain all bond lengths, including those involved in hydrogen bonding. The temperature was set to 300.0 K with 0.1 ps temperature-coupling. The “md” integrator was also used in the following 10.0 ns equilibration phase. The compressibility was set to 4.5 × 10^−5^ bar implementing the Berendsen pressure-coupling scheme. The same parameters associated with the peptides or the CNT in the complex mode were applied to simulate the individual free peptides (i.e., the P1’–P10’) or the unbound CNT (16,0). However, due to the systems’ smaller size, compared to the C1 and C2, the duration of the PR and equilibration steps were set to 2.0 ns and 4.0 ns, respectively. The conformation, obtained at the 10.0 ns time frame of the equilibration phase, was used as the starting conformation for the 1.500 μs MD data collection. The root mean square deviation (RMSD) of the complexes were calculated according to Maiorov *et al*.,^72^ for convergence assessment and estimating each conformation’s deviation from its corresponding reference structure throughout the MD trajectory.^73^ According to the RMSD data, the C1 and C2 systems were converged at ∼0.800 μs. The simulations were continued for extra 0.700 μs after the convergence to allow sufficient time for the peptides or the complexes to explore a broader range of conformational space. Thus, the data from 0.800–1.500 μs of the simulations were analyzed. All calculations were run on Graham, the High-Performance Computing Cluster of Compute Canada. (**Equation 1** and **Figure S1A–B**)

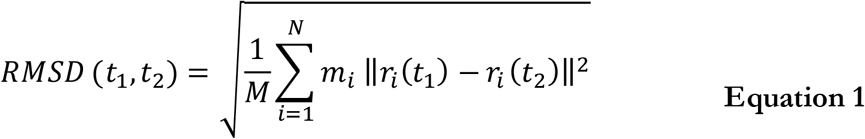

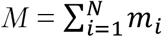 and *r*_*i*_ *(t)* are the position of atom *i* at *t* time.

### 2.3 Drugs’ Structure and Binding to Peptides

A ligand library was constructed utilizing SYBYL-X 2.1.1 software package (Certara Corporation©). The library consisted of seventeen (17) anticancer MT-targeting drugs: paclitaxel^37^, docetaxel^38^, taccalonolide^39^, epothilones (types A, B, D, and ixabepilone)^40, 41^, cyclostreptin^42^, dictyostatin^43^, discodermolide^44^, zampanolide^45^, laulimalide, peloruside A^46^, vinblastine^48, 49^, dolastatin^51^, phomopsin A^52^, and eribulin^53^. The structure of each drug was energy minimized using the steepest descent algorithm and the Tripos force field with an energy gradient ranging from 0.0005 kJ/mol to 1 kJ/mol for up to 10,000,000 iterations. For the docking of the ligands on the peptides, the representative conformations of the MD trajectories of the C1 and C2 systems were obtained through the RMSD-based linkage clustering method.^66^ The average structure of the most-populated conformation cluster was selected from the 0.800–1.500 μs time interval of each trajectory. For clustering, the conformation with the lowest gyration radius (R_g_ Å) was selected as the reference. FlexX^74-76^ docking software, embedded in v.2.1.8 of the LeadIT software package (BioSolveIT©), was used for the calculation and binding-energy-based (ΔG_binding_) ranking of the docking solutions of the drug-peptides conformations. The FlexX search algorithm is founded on a base-fragment and incremental construction. The interaction energies are calculated according to the Böhm scoring function^77^. Prior to the docking calculations, all the peptides’ amino acids were defined as potential targets for the drugs binding, allowing the algorithm to distinguish the best sites of interactions. The binding profiles of the drugs in the highest rank (i.e. the lowest ΔG_binding_) were inspected for their potential as cargo for loading on the CNT as a carrier.

## 3. Results And Discussion

The tubulin lateral-associated peptides (i.e., the P1 to P10) include amino acids with hydrophobic moieties that interact with the hydrophobic CNT surface through vdW forces. In contrast, their hydrophilic moieties make polar interactions with the solvent molecules or the adjacent peptides. The distance of 2.5–6.0 Å between an atom of the CNT and a peptide’s was a criterion for considering an inter-atomic interaction for calculating its Lenard Jones (LJ) potential interaction energy. ^78^ The peptides’ binding affinities to the CNT were affected by increasing their interaction frequency and the CNT-peptides solvent-accessible surface (SAS) that was calculated according to Eisenhaber *et al*.,^79^ implemented in the Gromacs package.^66, 67 80, 81^ (**Equation 2**)

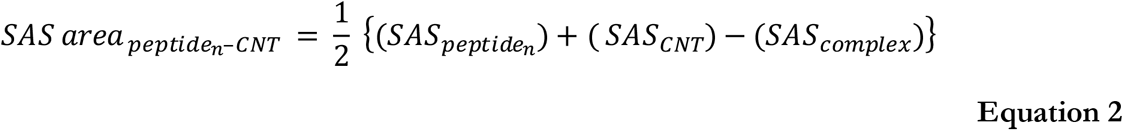

“n” represents a peptide index.

### 3.1 Analyses of the P1 to P10 Binding on the CNT

The analyses of the MD trajectories of the C1 and C2 systems showed that most of the peptides were bound to the CNT without any detachment from the tube surface; however, the P9 showed partial unbound conformations during 0.800–1.500 μs of the simulation. (**Figure 1C–D**)

A more detailed analysis of the observations, starting with the best binding peptides, was elucidated as follows:

#### 3.1.1. The P4 and the P9

The P4 (α-M-loop) from tubulin α-subunit possesses seventeen residues (Y_272_APVISAEKAYHEQLSV_288_), coded as 272–288 (1JFF.pdb^37^). It is equivalent to the P9 (β-M-loop) from β-tubulin, also with seventeen residues (P_274_LTSRGSQQYRALTVPE_290_), numbered as 274–290 in the α-β tubulin heterodimer structure in 1JFF^37^. The similarity between their sequences is 60.0% (identity of 26.7%). The P4 and P9 were simple loops in the tubulin heterodimer structure and mainly folded as coil and bend configurations when bound to the CNT during the MD simulation. The P4 showed a more effective binding pattern than the P9 presenting the lowest average distance to the CNT(0.30 nm), accounting for the most potent vdW interactions to the CNT, among the tubulin lateral segments (i.e. the P1–P10). The Leu286–Glu290 of the P9 moved away from the CNT (average distance > 0.57 nm) in favor of hydrogen bonding with the P6 hydrophilic amino acids and water molecules. However, its Pro274–Ala285 segment remained firmly bound to the CNT at a distance < ∼0.30 nm such that it made the P9 the second strongest CNT-binder among the P1–P10. (**Figure 2, Table S4** and **Table 4**)

**Table 4:** The average LJ energy, SAS area, interaction frequency number (at a distance ≤ 6.0 Å) between each peptide (i.e., the P1–P10) and the CNT over the simulation time, listed according to the binding strength to the CNT.

| Peptide<br>on the CNT | LJ energy<br>(kJ/mol) | SAS area (nm<sup>2</sup>) | Nº of<br>Interactions |
| --- | --- | --- | --- |
| **P4** (α-M-loop) | $-425.43 \pm 16.21$ | $6.05 \pm 0.28$ | $3,635 \pm 139$ |
| **P9** (β-M-loop) | $-375.84 \pm 27.29$ | $5.90 \pm 0.43$ | $3,373 \pm 228$ |
| **P6** (β-H1-B2) | $-314.47 \pm 12.18$ | $5.56 \pm 0.27$ | $2,823 \pm 113$ |
| **P2** (α-H2-B3) | $-245.41 \pm 8.92$ | $4.27 \pm 0.18$ | $2,304 \pm 83$ |
| **P7** (β-H2-B3) | $-238.11 \pm 10.47$ | $4.13 \pm 0.22$ | $2,046 \pm 96$ |
| **P10** (β-H10) | $-239.22 \pm 20.56$ | $4.40 \pm 0.37$ | $2,076 \pm 180$ |
| **P1** (α-H1-B2) | $-219.69 \pm 28.50$ | $3.64 \pm 0.32$ | $1,840 \pm 206$ |
| **P5** (α-H10) | $-172.03 \pm 10.03$ | $3.38 \pm 0.19$ | $1,783 \pm 88$ |
| **P8** (β-H4-T5) | $-104.40 \pm 12.33$ | $1.97 \pm 0.21$ | $865 \pm 101$ |
| **P3** (α-H4-T5) | $-101.51 \pm 16.58$ | $2.08 \pm 0.25$ | $975 \pm 147$ |

**Figure 2:**
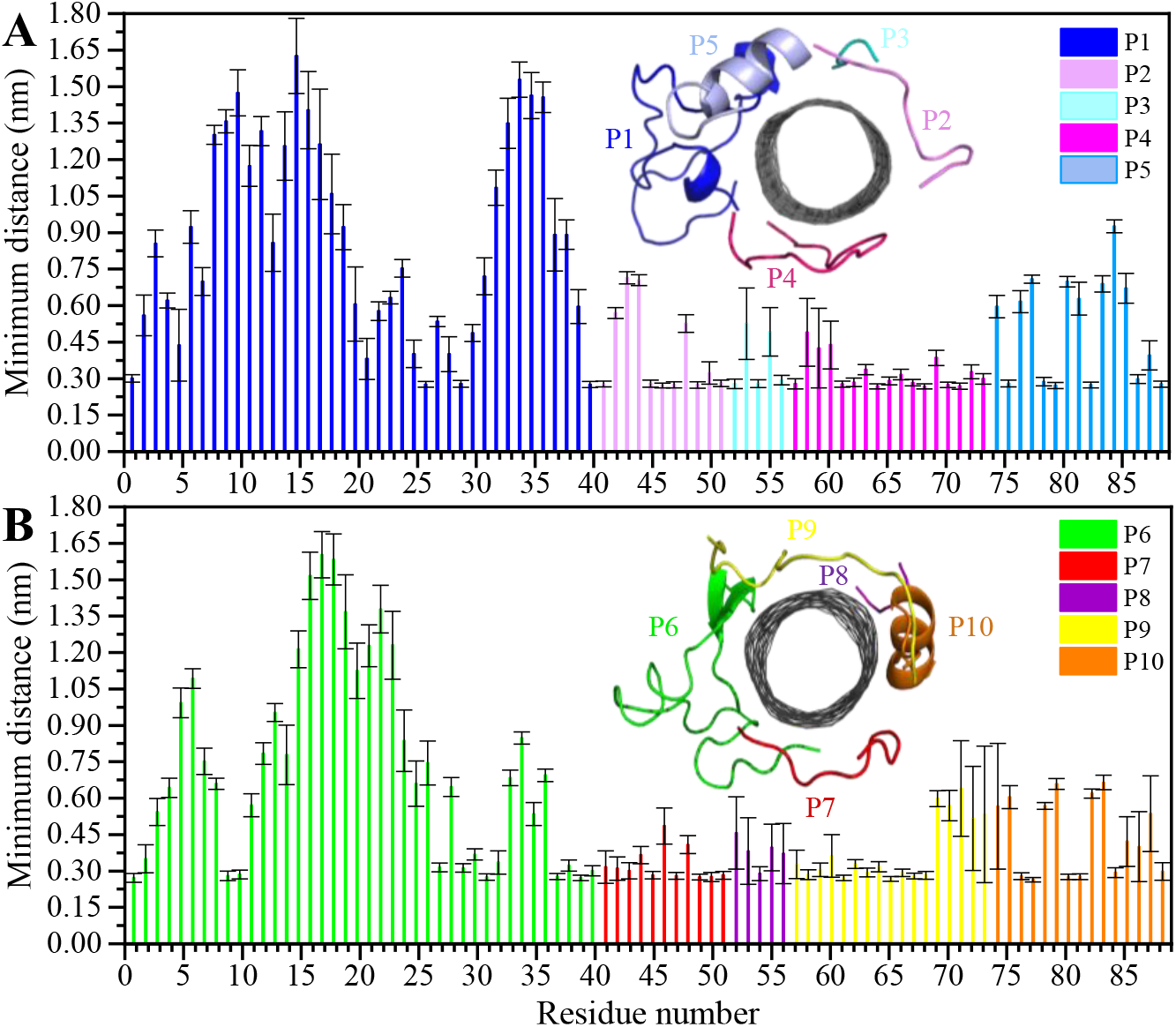
Residue-based binding analysis of the minimum distance between the centers-of-mass (COM) of the CNT and each of the: (**A**) P1 to P5 (the C1 system), and (**B**) P6 to P10 (the C2 system).

The P4 conformations in the bound state had the most effective binding affinity due to the peptide’s well-distributed side chains on the CNT, adapting satisfactorily to the tube’s curvature that favored maximum vdW interactions. The P4 binding affinity was reinforced by a combination of vdW and polar interactions through the nearest amino acids, possessing aromatic, cyclic, positively charged, and aliphatic moieties (Tyr282, Tyr272, Gln285, Ile276, Lys280, His283, Pro274, Ala281, and Val275). The most extreme fluctuation of the P4 was during 1.160–1.190 μs, caused by its Tyr272 movements. It rotated from its perpendicular to parallel position to the CNT as it was stimulated by a water molecule that bridged a hydrogen bond with the tyrosine hydroxyl moiety and an oxygen atom of Glu284. That caused a steric hindrance, triggering Pro274 desorption from the CNT, and consequently weakened the LJ potential energy of the P4-CNT by ∼27 kJ/mol. That occurred along with the increase of the distance of their centers of mass (DCOM) for ∼1 Å. Tyr272 at the N-terminal caused significant instability and weakened the vdW interactions of the P4 with the tube during 45% of the simulation time. Despite that, the P4 was identified as the best CNT-binder among the P1–P10. The observations suggest that Tyr272 could be replaced with Phe to eliminate Tyr hydroxyl moiety and its H-bonding property. (**Figure 3**)

**Figure 3:**
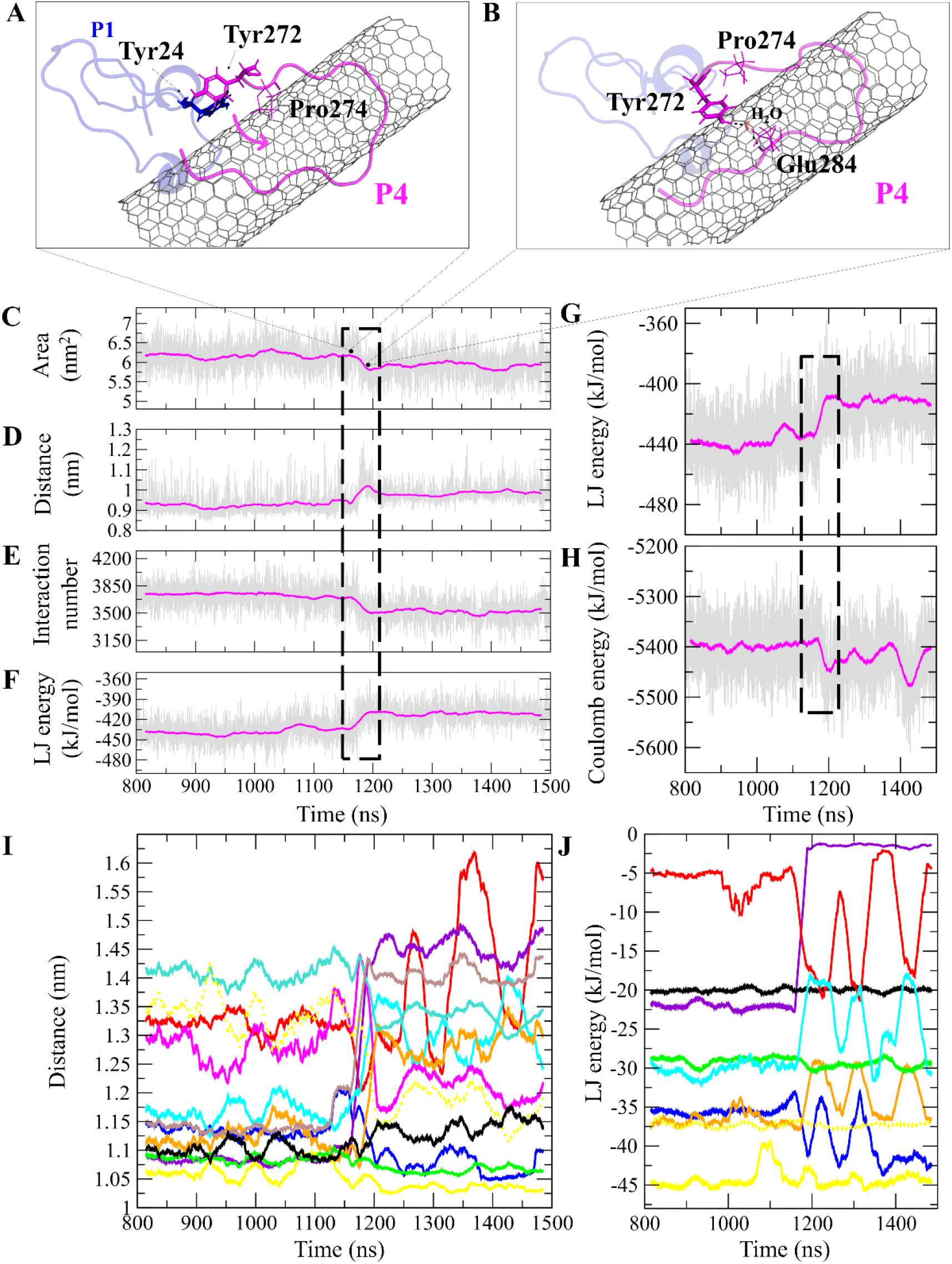
The P1 and P4, (**A**) and (**B**) movement of Tyr272 ring of the P4 from vdW interaction to Tyr24 of the P1. A water molecule bridges Glu284 of the P4 to Tyr24, changing its aromatic ring position from perpendicular to parallel orientation on the CNT. The time evolution of properties between the CNT and the P4 (α-M loop): (**C**) SAS area, (**D**) DCOM, (**E**) interaction frequency (< 6.0 Å), and (**F**) LJ energy. The P4 intramolecular (**G**) LJ energy and (**H**) coulomb energy. (**I**) The average DCOM between the CNT and the P4 residues Tyr272 (blue), Ala273 (brown), Pro274 (violet), Val275 (red), Ile276 (orange), Lys280 (cyan), Ala281 (black), Tyr282 (yellow), His283 (green), Glu284 (turquoise), Gln285 (dotted yellow), and Leu286 (magenta). (**J**) The average LJ energy of the CNT with: Tyr282 (yellow), Tyr272 (blue), Gln285 (dotted yellow), Ile276 (orange), His283 (green), Ala281 (black), Pro274 (violet), Val275 (red), and Lys280 (cyan). Graphs were generated with averaging every 300 frames of the 1.500 μs MD trajectory.

The closely interacting P9 residues with the CNT were either positively charged or consisted of an aromatic moiety (Arg278, Arg284, Tyr283). Despite its Leu286, Val288, and Pro289 being intrinsically hydrophobic, their associated segment in the P9 (Leu286–Glu290) was hindered from firmly binding to the tube due to the more dominant effect of the hydrophilic fragments and their hydrogen bonding with water molecules, or the electrostatic interaction of Glu290 with Arg48 of its neighboring peptide, the P6. They, together, caused desorption of the P9 segment from the CNT during 0.864–0.871 μs and 1.027–1.073 μs time intervals. However, the overall P9’s LJ energy improved over time due to the more frequent conformational changes of the Leu286–Glu290 segment favoring the CNT binding. The P9 instability and its nearly half-sequence desorption can be abolished using a CNT with a greater diameter or length than the CNT (16,0), capable of accommodating a broader range of the P9 conformations. In addition, replacing the P9’s Glu290 with a hydrophobic residue can eliminate its electrostatic interactions with the P6 and the P1 and improve its binding to the CNT. (**Figure 4B–C** and **Figure S2**)

**Figure 4:**
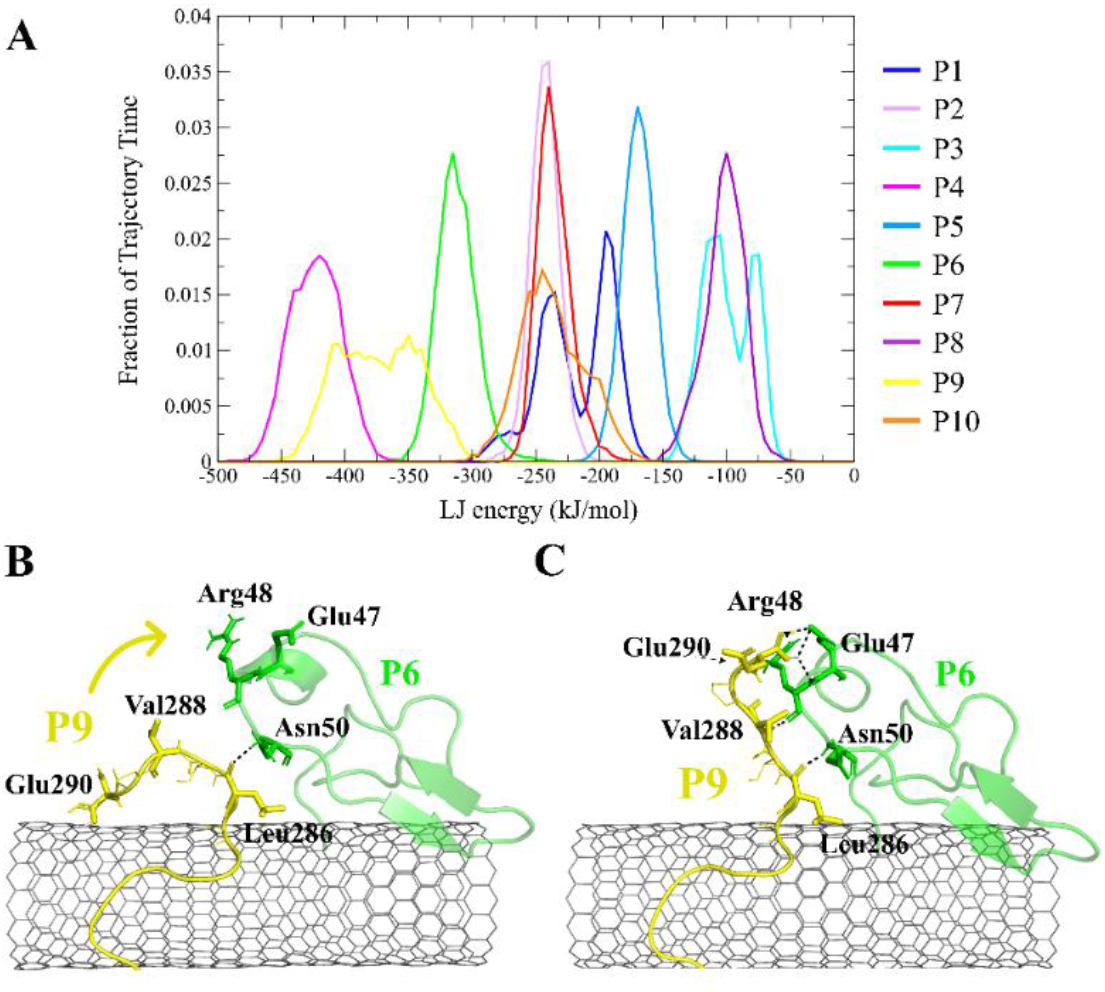
(**A**) The fraction of the simulation time (after convergence at ∼0.800 μs) that each peptide was bound to the CNT versus their LJ potential energy (vdW interaction). The representative conformations of the P9 (Leu286–Glu290) fragment (**B**) binding to the CNT, (**C**) distancing from the CNT to H-bond to the P6 at the Glu47–Ans50 portion. The dashed lines represent hydrogen bonds.

The P9 and P10 had the shortest CNT-binding lifetime, ∼10% and ∼17% of the simulation time after the convergence (i.e. 0.800 μs). The broadest LJ potential energy distribution curves as in the P4 (∼−360–−480 kJ/mol) or the P9 (∼−300–−500 kJ/mol) indicated a highly stable hydrophobic binding to the CNT. In contrast, the P2 had ∼27%, the P7 ∼33%, and P5 ∼32% dwell time on the tube interacting to the CNT more frequently than the other peptides (i.e., the P4 ∼19%; the P9 ∼10%). The long-lasting binders (the P2, P5, and P7) remained at the tube’s central region involving their adjacent peptides, which limited their conformational freedom. (**Figure 4A**)

#### 3.1.2. The P1 and the P6

The P6 is another MT-lateral segment, a simple loop that links the H1 helix and the B2 strand of β-tubulin. It consists of forty (40) residues (I_24_SDEHGIDPTGSYHGDSD-LQLERINVYYNEATGNKYVPRA_65_) coded 24–65 in 1JFF.pdb^37^. The P6 was the third strongest binder to the CNT among the P1-P10. The P6 most considerable contribution to its LJ potential energy was made by Arg64 and the phenyl ring of Tyr52. The P6’s Arg64 was stabilized at a ∼3.5 Å distance to the CNT due to its H-bonding to Asp90 in the P7. The binding energy of Tyr52 fluctuated as its phenyl ring rotated from parallel to the tube from 0–45°. The intra-peptide H-bond of Tyr52 to Asn54 also weakened the peptide’s binding to the CNT, suggesting that the Asn mutation with a more hydrophobic amino acid (e.g. Leu) prevents the disadvantageous interaction. The LJ energy was favorably affected by water molecules’ H-bonds to the neighboring residues of Tyr52 during 1.182–1.219 μs, causing the Tyr ring to adapt a parallel conformation to the CNT that was also supported by the H-bond with the Arg284 of the P9 and consequently strengthened the vdW force between the P6 and the CNT. Similarly, a Tyr and an Arg interacted in the P2, P5, P6, P9 (each consisting of the Arg), and the P2, P4, and P9 (each consisting of the Tyr), resulting in the improvement of the binding conformation of the latter group to the CNT, which indicates the importance of the Tyr for the CNT coating (**Table 4** and **Figure S3**)

The P1, from α-tubulin, is known as the α-H1-B2 loop in the α, β-heterodimer tubulin, with forty (40) residues (Y_24_CLEHGIQPDGQMPSDKTIGGGDDSFNTFFSET-GAGKHVP_63_), coded as 24–63.^37^ Due to the dense intra-peptide interactions in the P1, only a few hydrophobic residues consisting of aromatic or cyclic moieties (i.e., Phe49, Tyr24, Phe52, Pro63, His28, and Gly44) were available for binding to the CNT. The P6 is the equivalent peptide to the P1 in β-tubulin. They have 75.7% sequence similarity and 32.4% identity. The former covers a broader tube surface area than the latter. Compared to the P1’, the P1 binding to the CNT resulted in a helix folding (i.e., α- and 3-helix) at Cys25–His28 and Asp47–Asn50 segments that demonstrated the impact of the CNT binding. That was also shown in the P6 compared to the P6’, as it maintained its β-strand configuration at Tyr52–Glu55 and Asn59–Val62, similar to its secondary structure in the MT. The P6’ highly fluctuated in a coil configuration since it was influenced by its exposure to the aqueous environment. (**Figure 5A-F** and **Table S4**)

**Figure 5:**
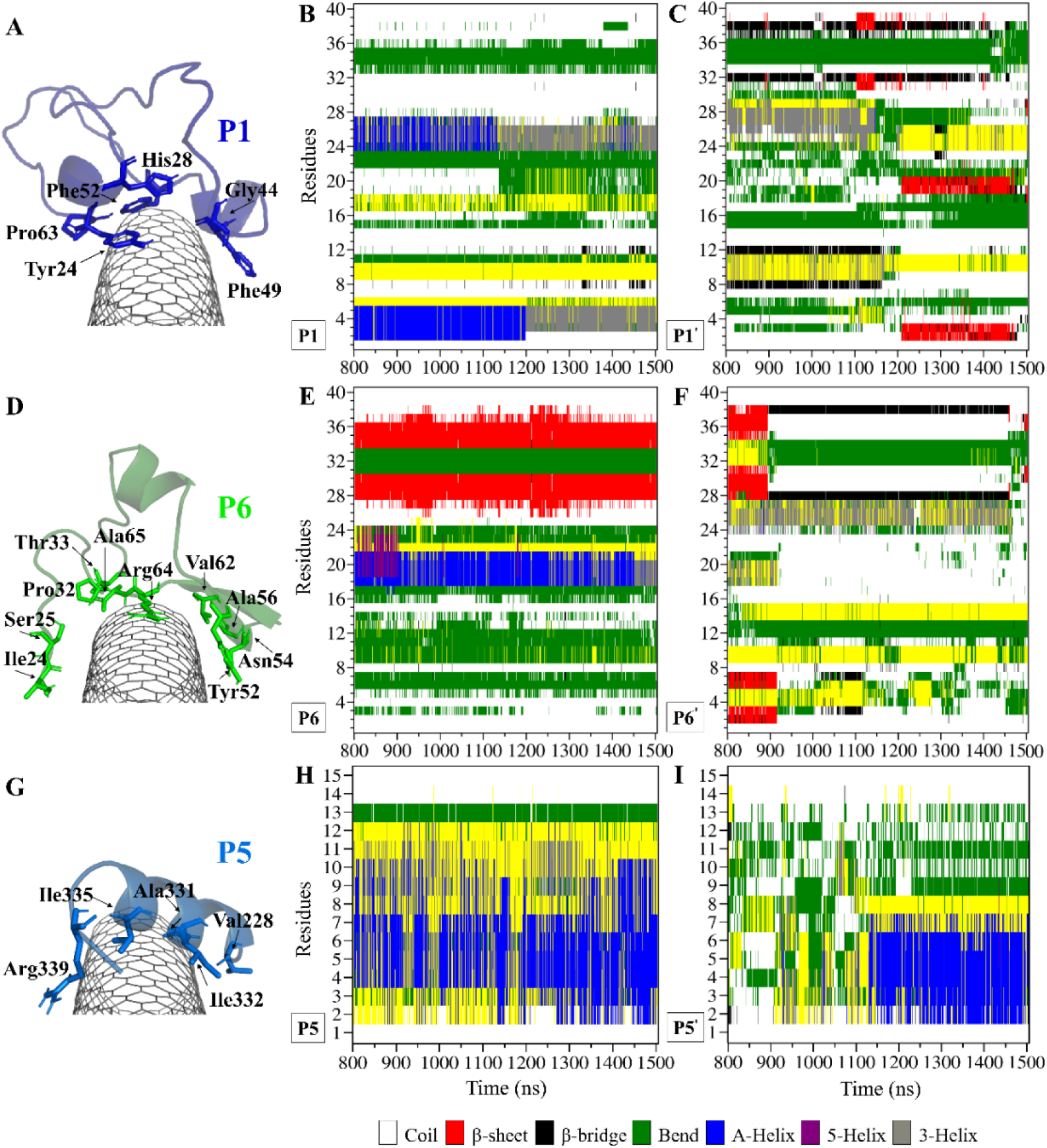
Binding profile of (**A**) the P1 (the C1 system) with a hydrophobic cyclic residue moiety structure core, (**D**) the P6 (the C2 system) coating a wider CNT surface area than the P1, and (**G**) the P5 (the C1 system). The secondary structure (SS) analysis is based on the Dictionary Secondary Structure of Protein (DSSP) algorithm during 0.800–1.500 μs of the (**B**) P1, (**E**) P6, and (**H**) P5 on the CNT, and their CNT-peptide-free states the (**C**) P1’, (**F**) P6’, and (**I**) P5’. The residue numbering of 1–40 corresponds to Tyr24–Pro63 in the P1 and Ile24–Ala65 in the P6.

There were three major conformational changes in the P1 that occurred during *i*. 1.130–1.155 μs, *ii*.1.375–1.380 μs, and *iii*. 1.430–1.435 μs. Unlike the third time interval, during the first two, the P1-CNT binding improved due to His28 and Gly44 movements and hydrophobic binding to the CNT. However, the third was related to Tyr24 and its ring’s rotation from parallel, in a range of 0°–30° with respect to the CNT’s long axis. Its hydroxyl moiety made H-bond to water molecules and formed H-bond bridges to His28 and Glu27. Those water molecules pulled the ring away from the CNT that weakened the vdW forces. That suggests a mutation at Tyr24 with Phe will omit its hydroxyl H-bonding effects and prevent declining P1 affinity to the CNT. (**Figure S4**)

Comparing RMSD of the P1 and the P6 to their respective CNT-free conformations, the P1’ and the P6’, demonstrates the CNT’s role in refining their secondary structure with steady binding ability. (**Figure S1C–D** and **Figure 5A-F**)

#### 3.1.3. The P2 and the P7

The P2, from α-tubulin, has eleven residues (R_79_TGTYRQLFHP_89_) with amino acids coded as 79–89 in 1JFF.pdb^37^, known as the α-H2-B3 loop. It is equivalent to the P7 of β-tubulin, also with eleven residues (G_81_PFGQIFRPDN_91_), coded as 81–91 at the β-H2-B3 loop^37^. The P2 and P7 have 77.8 % sequence similarity and 44.4% identity. They adapted a combination of folding as coil, bend, and turn during the MD simulation time, distinct from the lengthier peptides (e.g., the P1 and P6) at the bound state on the CNT. (**Table S4** and **Figure S5A–B**)

The turn formation prevented the maximum LJ interaction of the P2 with the CNT at the Thr80–Thr82 segment, caused by Tyr83 spatial hindrance and the frequent perpendicularly reorienting Tyr83 and Phe87 aromatic rings. The P2’s N-terminal (Arg79– Tyr83) wrapped around the CNT in a crook shape, creating a condition for a steady intra-peptide H-bond between Arg79 and Arg84. That facilitated a stable flat conformation of Arg84 aliphatic moiety and π–π interactions of its guanidine moiety to the CNT. The Arg, along with the contribution of the bridging water molecules, formed H-bond to Gln85. The N-terminal’s Arg79 was also frequently H-bonded to Tyr83, which increased the distance between the Thr80–Tyr83 segment and the CNT. Replacing Tyr83 and Phe87 of the P2 with a simpler hydrophobic residue (e.g. Leu) could reduce their steric hindrance, result in better exposure of the P2 to the CNT, and increase the P2’s binding affinity. In addition, mutating Arg84 also with a more hydrophobic amino acid can reduce the potential H-bond formation. In the P7, Phe87 and Phe83 rings were also perpendicular to the CNT, similar to Phe87 of the P2. Their conformations prevented the P7 full interactions, negatively affecting the P7-CNT binding and weakening the LJ energy. As suggested for the P2, replacing Phe83 with a non-aromatic hydrophobic residue is expected to enhance the P7’s binding property. (**Figure S6–S7**)

#### 3.1.4. The P3 and the P8

The P3 and P8 were the shortest sequences among the P1-P10 with low intrinsic hydrophobicity. The P3 is the H4-T5 loop in α-tubulin, with five amino acids (L_157_SVDY_161_), numbered 157–161.^37^ The five residues shaping the P8 are from β-tubulin (E_159_EYPD_163_), coded as 159–163.^37^ The P3 and the P8 showed supporting roles for their neighboring peptides. For instance, the inter-peptide interaction profile of the P3 showed that it mainly involved the hydrophobic, polar, and charged amino acids of the P2 and P5, demonstrating its role was primarily auxiliary to the conformation of its adjacent peptides’ to CNT-binding. (**Table 4** and **Figure 6**)

**Figure 6:**
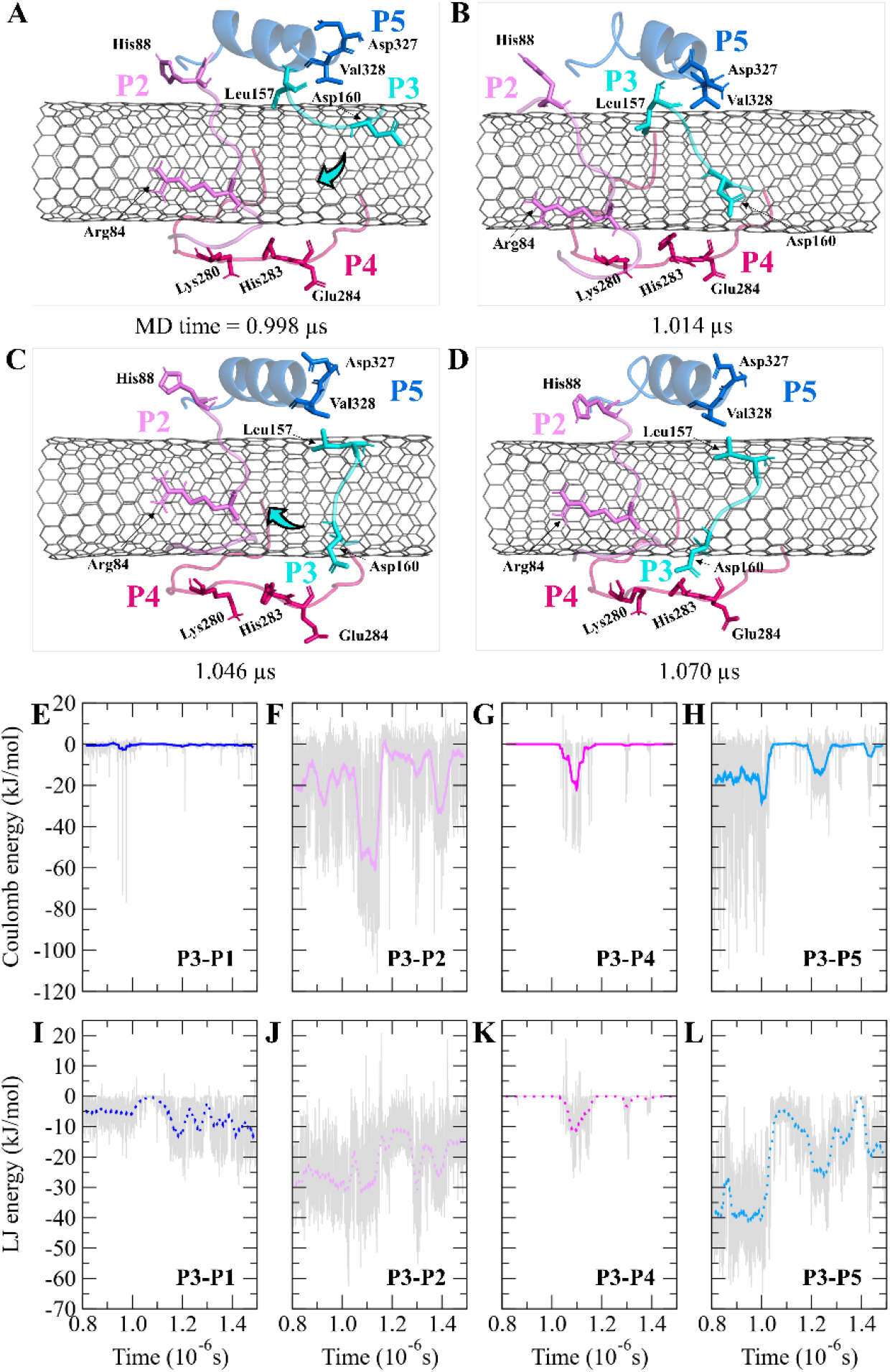
The P2–P5 binding to the CNT. Inter-peptide coulomb and LJ energy of (**A, E**) the P1-P3, (**B, F**) the P2-P3, (**C, G**) the P3-P4, and (**D, H**) the P3-P5 on the CNT. Graphs were generated with averaging every 300 frames.

#### 3.1.5. The P5 and the P10

The P5 belongs to α-tubulin and comprises fifteen (15) amino acids (D_327_VNAAIAT-IKTKRSI_341_) coded as 327–341 in 1JFF.pdb.^37^ Unlike the other peptides, the P5 lacks aromatic or cyclic side chains, affecting the magnitude of its LJ potential energy in the CNT binding. The P10 of β-tubulin also has fifteen amino acids (D_329_EQMLNVQNKNSSYF_343_), equivalent to 329–3431.^37^ The sequence similarity and identity between the P5 and P10 are 66.7% and 16.7%, respectively. Due to the CNT presence, the P5 and the P10 folded into α-helix, 3-helix, and turn configurations during the MD simulation time. The unstable turn configuration of the P5 was caused by Arg339. It shifted away from the tube, disturbed the intra-peptide H-bonding of the P5 that is necessary for maintaining the helicity. Arg also caused steric-hindrance to other amino acids’ binding to the CNT that cost ∼19 kJ/mol in LJ potential energy of the P5-CNT binding. Mutating Arg339 with serine could eliminate the Arg size-related steric hindrance while contributing to the helix forming H-bonds. (**Figure 5G-I** and **Figure S8–S9**)

#### 3.2. Peptides’ Stability on the CNT and the Complex Solubility

The residues’ root mean square fluctuation (RMSF) data were inspected to evaluate the peptides’ secondary structure compared to that in the MT or the peptides in their CNT-free states (i.e., the P1’–P10’). A residue with an RMSF ≤ 2.0 Å was considered steady; otherwise, it was classified as highly flexible. According to that criterion, the peptides from β-tubulin, frequently appearing with random coil configuration and consisting of polar or ionic residues, presented higher RMSF than their counterparts in α-tubulin (i.e., the P1–P5). Comparing residues’ RMSFs in the CNT-bound peptides with their free forms showed the configurational stability of the former, as seen in the C-terminal of the P1, the P2, and the P4–P7. (**Figure 7** and **Table S5**)

**Figure 7:**
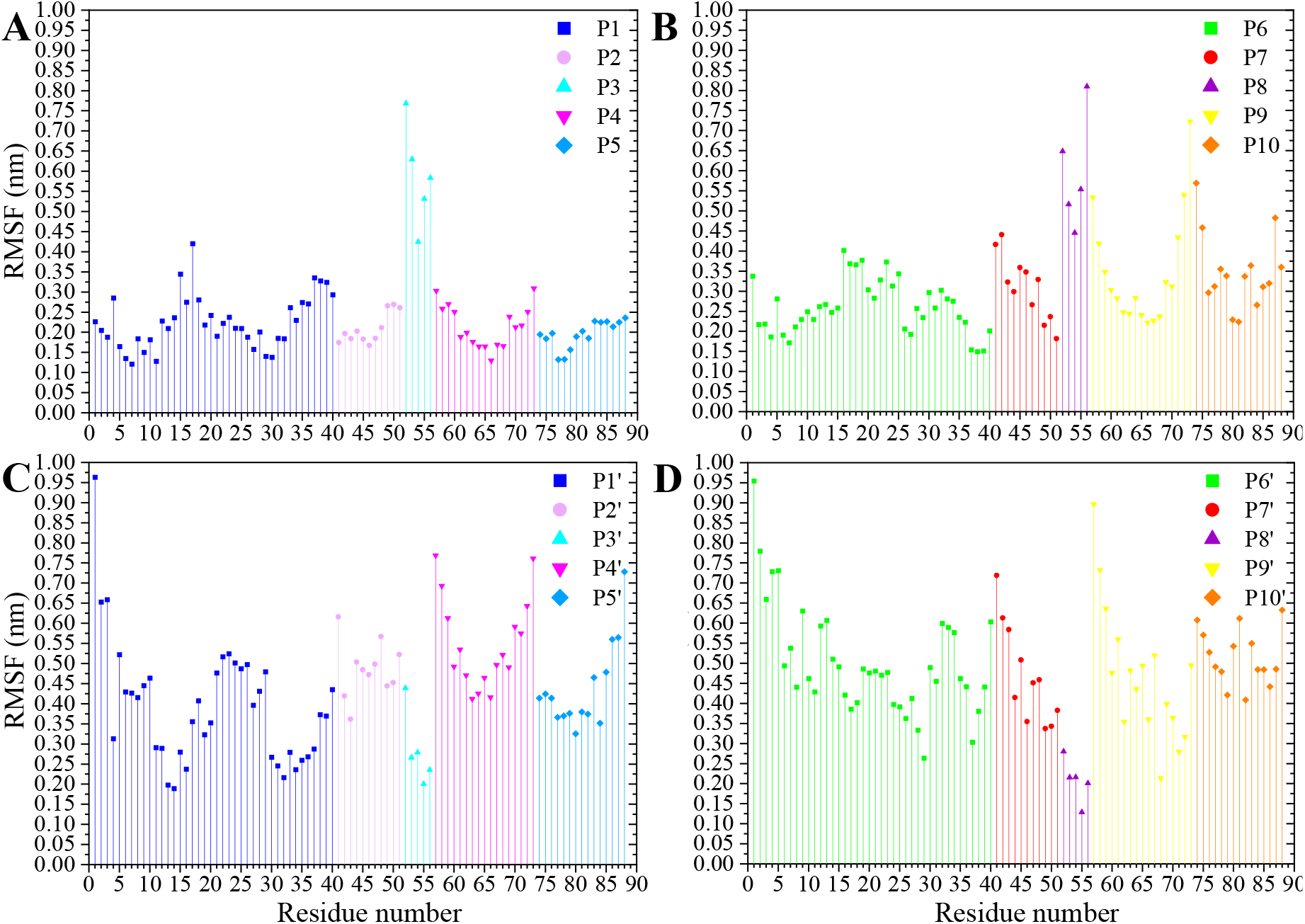
The RMSF of the amino acids (**A**) the P1–P5 and (**B**) the P6–P10 on the CNT and their CNT-free structure, (**C**) the P1’–P5’ and (**D**) the P6’–P10’, during 1.500 μs MD trajectory.

A ∼3 Å dehydration gap near the CNT wall at its exterior and interior regions was due to the hydrophobicity of the pristine CNT that functioned in favor of the vdW interactions to the peptides’ hydrophobic side chains and the CNT-peptides LJ energy. (**Figure S10**) The water molecules triggered the charged and polar residues of the CNT-bound peptides to reorient towards the solvent and, consequently, altered the peptides’ polar surface distribution on the tube. The peptides bound to the tube via polar or positively charged amino acids simultaneously formed H-bonds with the aqueous solvent molecules; for instance, those in the P1 (Glu27, Asp33, Lys40, Asp47, Lys60, and His61), the P4 (Glu284), the P5 (Asp327 and Lys336), the P6 (Asp26, His28, Asp39, Asp41, and Arg48), the P7 (Arg88), the P9 (Glu290), and the P10 (Asp329 and Lys338). That suggests their potential to improve CNT solubilization and dispersibility in water. The peptides’ effects on the CNT’s solubility vary depending on the number of H-bonds formed. For instance, the P6 had a considerably higher number of H-bonds with its surrounding water molecules (in the C2) than the P1 (in the C1). The P2, P8, P9, and P10 displayed a superior potential for influencing the CNT solubility over the P3, P4, P5, and P7. (**Figure 8, Figure S11**, and **Table S6**)

**Figure 8:**
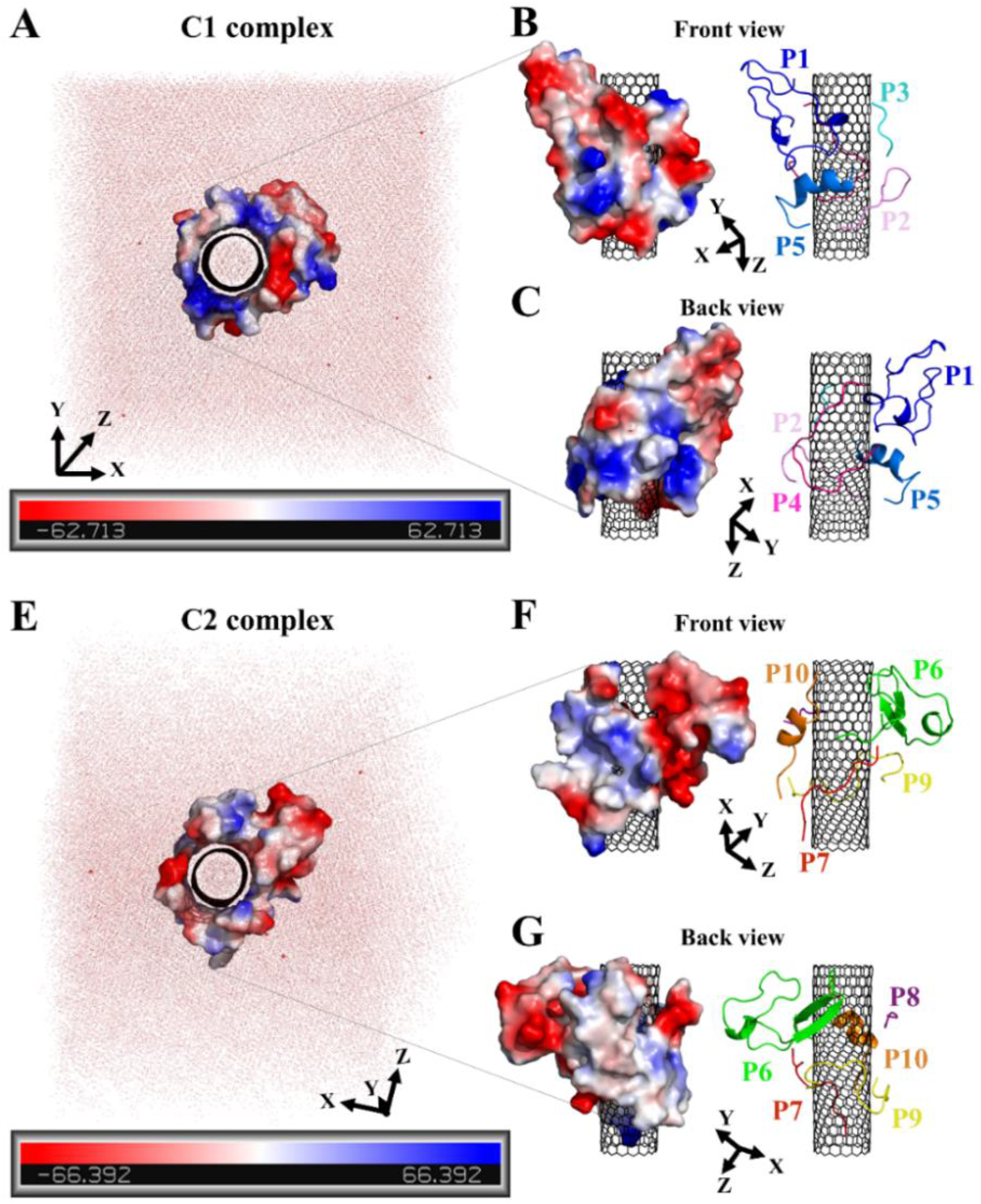
The peptides’ polar surface distribution in (**A**) the C1 and (**E**) the C2. The front and back views of (**B, C**) the C1 and (**F, G**) the C2. Negative charge or polarity (red), and the positive charge or polarity of the amino acids (blue).

The hydrophilic surface of the β-tubulin peptides (the C2) was ∼51.5 nm^2^, which was more significant than of the peptides from α-tubulin (the C1), ∼44 nm^2^ since the C2 peptides have 54 polar side chains (including the charged residues) compared to 48 of the C1. Thus, the C2 system functioned better than the C1 in favor of CNT’s solubility. Conversely, the peptides from α-tubulin, hydrophobically interacting with the CNT, resulted in a higher coating in the C1 of ∼22 nm^2^ compared to ∼19 nm^2^ surface area in the C2. (**Table S7**)

The CNTs randomly and diffusively traversed from their initial position in the solvent box of the peptide-bound and unbound systems. The CNT’s radius of gyration, R_g_^66, 67^, was calculated according to Equation 3. The R_g_ of the unbound CNT was less than its peptide-bound form, indicating the broad distribution of the peptides’ atoms around the tube’s center of mass in the C1 and C2 during the complexes’ rotational and transitional motions. It was a consequence of the peptides’ hydrophilic nature and their interactions to the aqueous polar environment, causing the hydrodynamic shell around the CNT-peptides complexes to expand and increase the R_g_. (**Equation 3** and **Figure 9**)

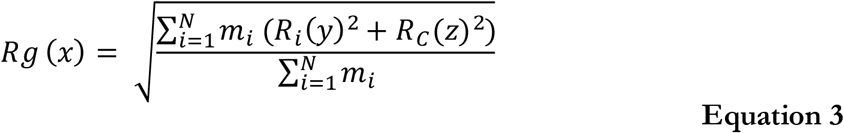

**Figure 9:**
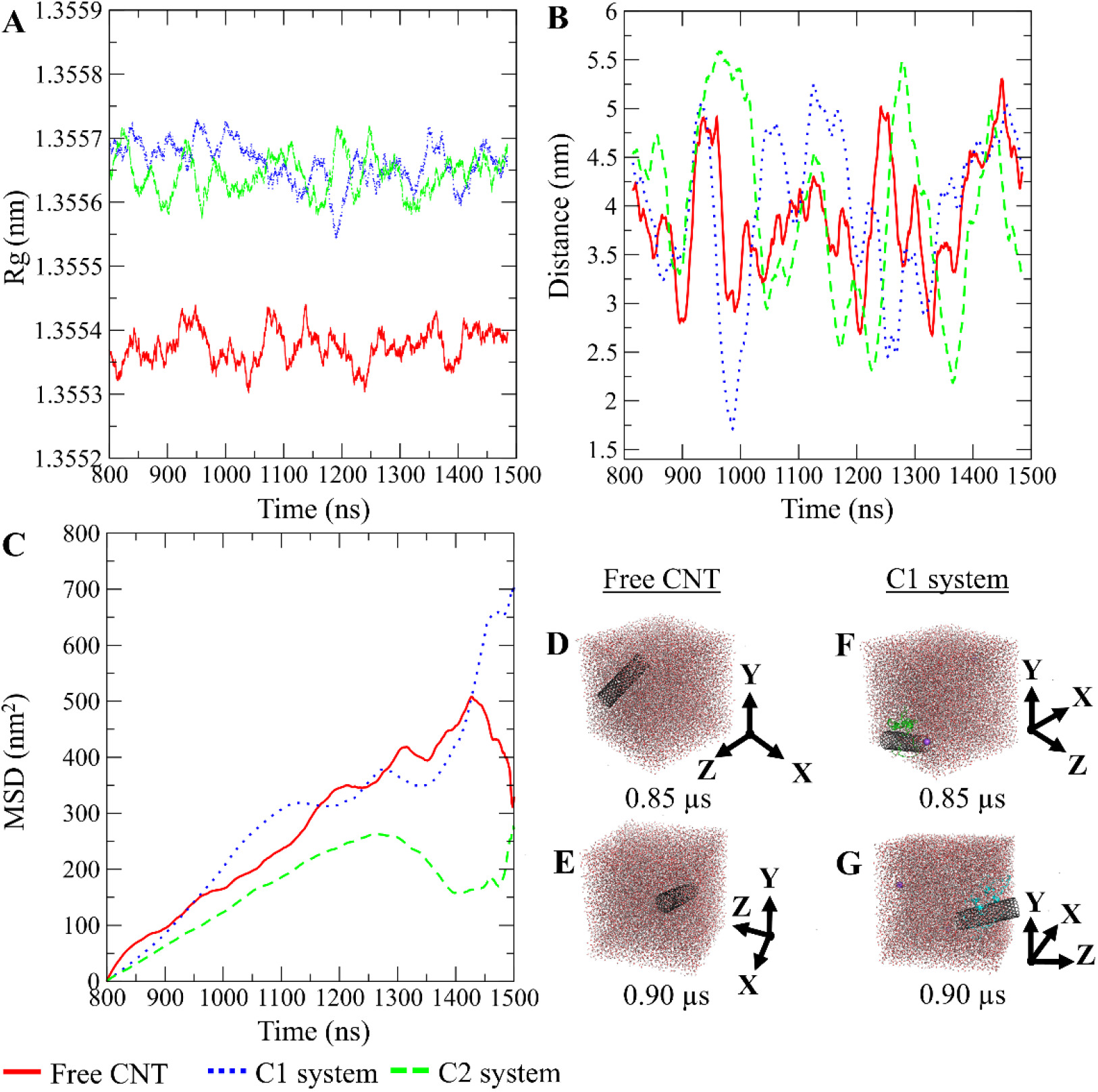
The pristine CNT, the C1, and the C2 systems. (**A**) R_g_, (**B**) the DCOM between the water box and each CNT, (**C**) Mean Squared Displacement (MSD). (**D, E**) The pristine CNT and (**F, G**) the C1 system (the P1–P5) simulation boxes. Graphs were generated with averaging every 300 frames.

Radius of gyration of a molecule (m) about the x, y, and z axes.

The CNT’s mean squared displacement (MSD)^82^ was calculated as a function of time. The diffusion coefficient (D_A_) was also calculated, according to Einstein’s relation^83^, to calculate the CNT’s position deviation. (**Equation 4–6**)

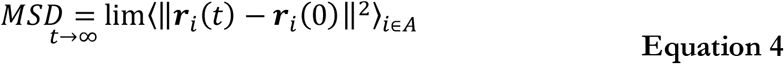

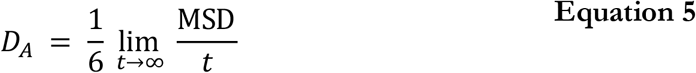

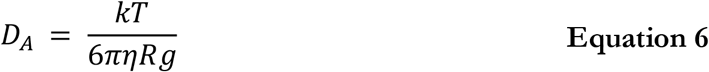

*r*_*i*_ (*t*) *− r*_*i*_ (0) is the distance (vector) traveled by the center-of-mass (COM) of an *i*^th^ particle over time *t* in a solvent with *η* viscosity, *k*, Boltzmann constant, and temperature T.

The D_A_ of the unbound pristine, the CNT-peptides in the C1, and the C2 systems were 1.21E+8 nm^2^/s, 1.38E+8 nm^2^/s, and 0.99E+8 nm^2^/s. They were obtained by least-squares fitting the best straight line of each MSD graph. The D_A_ estimated a restricted flux of the CNT-peptides complex in the C2 system, pertinent to its controlled motions by the more hydrophilic peptides of β-tubulin (the P6–P10) than α-tubulin’s (the P1–P5). The polar amino acids created a hydrodynamic shell around the hydrophobic CNT that carried several water molecules involved in the H-bond networks or polar interactions and thus directly affected the complexes’ R_g_. (**Table 5** and **Table S7**)

**Table 5:**
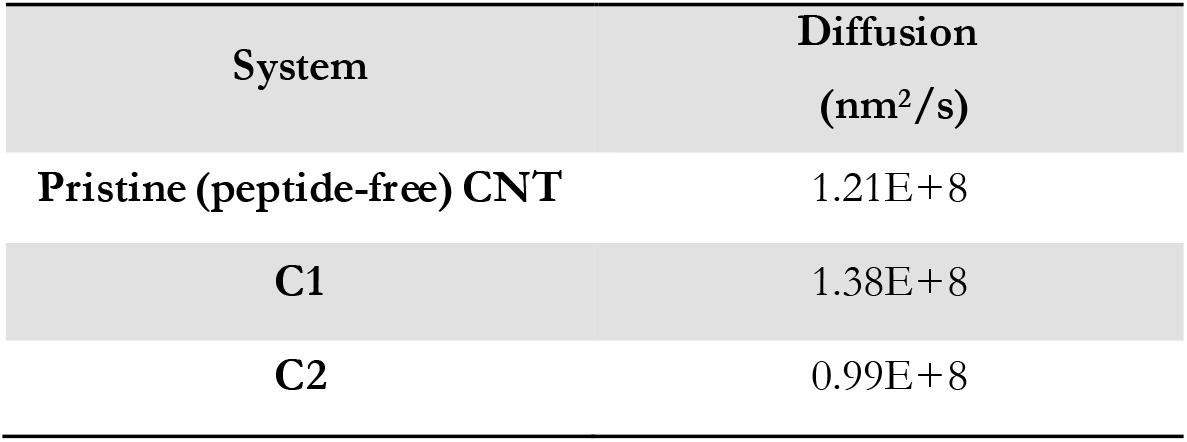
Diffusion coefficient according to the MSD data’s linearity trend.

| System | Diffusion<br>(nm<sup>2</sup>/s) |
| --- | --- |
| Pristine (peptide-free) CNT | 1.21E+8 |
| C1 | 1.38E+8 |
| C2 | 0.99E+8 |

The R_g_ determines the D_A_ and is influenced by several factors, including the peptides’ amino acid composition. The magnitude of the amino acids’ hydrophobicity or hydrophilicity regulates binding strength to the tube and the peptides’ tendency to stretch out towards the aqueous environment. Accordingly, they shrink or expand the hydrodynamic shell around the CNT and alter the R_g_ in inverse proportion to D_A_ as demonstrated by the correlation of the MSD and R_g_ data. (**Figure 9**)

### 3.3. Antimitotics Binding

The peptides’ binding to drugs was assessed using a molecular docking method. Seventeen MT-binding agents were studied that included paclitaxel (PTX)^37^, docetaxel^38^, taccalonolide^39^, epothilones (i.e., A, B, D, and ixabepilone)^40, 41^, cyclostreptin^42^, dictyostatin^43^, discodermolide^44^, zampanolide^45^, and dolastatin.^51^ The crystal structures of the drugs bound to α-β tubulin heterodimer exhibit they all are bound to at least one amino acid in the M-loop of β-tubulin. It also shows that the P5 (α-H10), the P6 (β-H1-B2), and particularly the P9 (β-M-loop) to be critical for the CNT functionalization for drug delivery due to their key roles in the binding of several ligands to the tubulin subunits. For instance, epothilone A (4O4I.pdb^46^) encounters the highest number of amino acids in the MT binding site at the β-M-loop (the P9) with Pro274, Leu275, Thr276, Arg278, Gln281, Gln282, and Arg284. Taxol (1JFF.pdb)^37^ binds to more than one protein segment, β-H1-B2 (the P6) and β-M-loop (the P9), similar to dolastatin (4X1I.pdb^51^) that is hosted in the MT by α-H10 (the P5) and β-M-loop (the P9) or phomopsin A (3DU7.pdb^52^) by α-H10 (the P5) and β-H1-B2 (the P6). (**Table 6** and **Figure 10**)

**Table 6:**
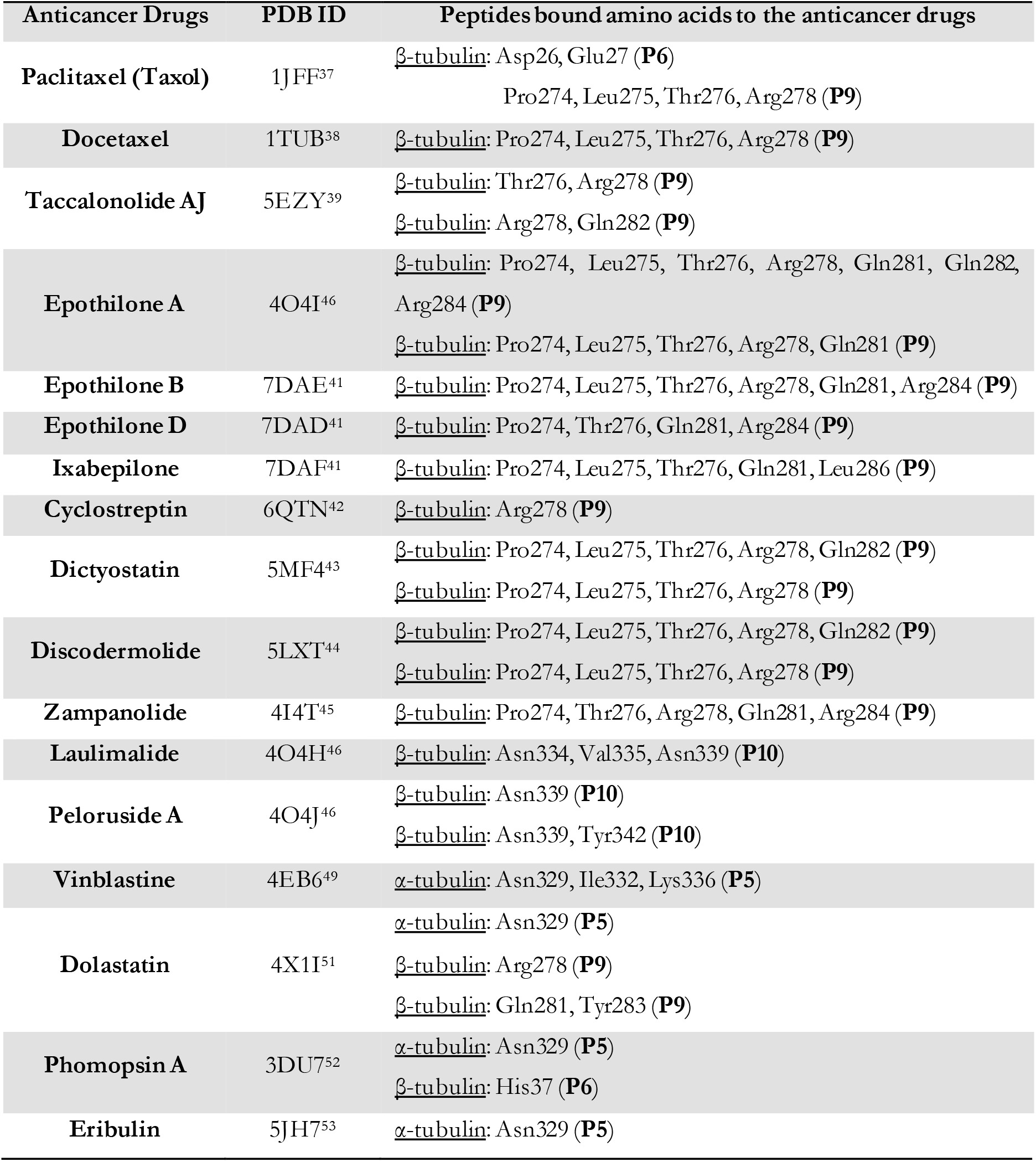
Microtubule targeting antimitotics and the crystal structure of their complex with α, β-heterodimer along with their identification code in the RCSB Protein Data Bank. The “α” and “β” refer to the tubulin α and β-subunits.

**Figure 10:**
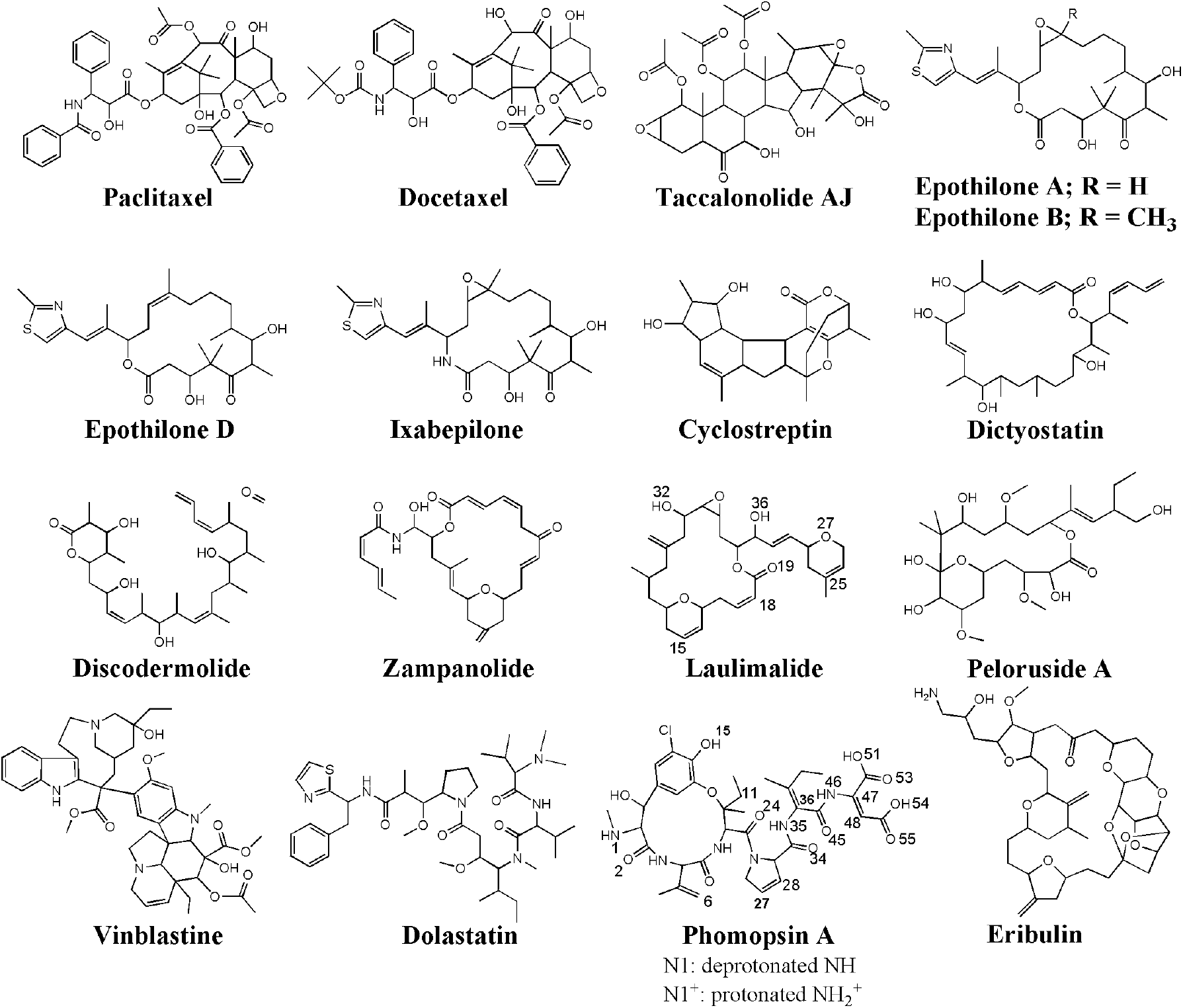
The chemical structures of the seventeen (17) microtubule targeting anticancer drugs in the ligand library, drawn and numbered using Chemdraw software.

According to the docking results, the two lowest binding energies of the ligand– peptides associated with phomopsin A (neutral and protonated at N1 atom) and laulimalide bound to the P6 and the P7 (in the C2). The neutral phomopsin A presented the highest binding affinity (−26.56 kJ/mol) among the ligands. (**Table 7** and **Figure 10–11**)

**Table 7:** The top 13 docking solutions of the anticancer tubulin ligands in the C1 and C2 complexes; ranked based on the binding energy.

| Agent | System | Binding energy (kJ/mol) | Ligand-binding peptides | Involving amino acids |
| --- | --- | --- | --- | --- |
| Phomopsin A (neutral) | C2 | -26.56 | P6 and P7 | <u>H-bond</u>: Glu27, His28, Leu42, Asn50, Arg64 (**P6**)<br><u>vdW</u>: Asp26, His28, Gly29, Arg64 (**P6**) Arg88 (**P7**) |
| Phomopsin A (protonated) | C2 | -17.99 | P6 and P7 | <u>H-bond</u>: His28, Gln43 (**P6**) and Arg88, Asp90 (**P7**)<br><u>vdW</u>: Asp26, Glu27, Gln43, Leu42, Asn91 (**P6**)<br><u>Ionic</u>: Glu290 (**P9**) |
| Laulimalide | C2 | -17.79 | P6 and P7 | <u>H-bond</u>: Glu27, Gln43, Asn50 (**P6**) Asp90 (**P7**)<br><u>vdW</u>: Glu27, His28, Gly29, Ile49 (**P6**) |
| Epothilone A | C2 | -17.01 | P6 and P7 | <u>H-bond</u>: Asp26, Glu27, His28, Gly29, Gln43 (**P6**) Arg88 (**P7**)<br><u>vdW</u>: His28, Arg48 (**P6**) |
| Epothilone D | C2 | -14.79 | P6 and P7 | <u>H-bond</u>: Glu27, Gln43 (**P6**) Asn91 (**P7**)<br><u>vdW</u>: His28 (**P6**), Arg88 (**P7**) |
| Cydostreptin | C2 | -13.75 | P7 and P10 | <u>H-bond</u>: Gln85, Ile86 (**P7**) Ser341 (**P10**)<br><u>vdW</u>: Gln85, Ile86 (**P7**) |
| Epothilone A | C1 | -12.06 | P1 | <u>H-bond</u>: Ile42, Gly44, Gly45 (**P1**)<br><u>vdW</u>: Asp47 (**P1**) |
| Discodermolide | C2 | -12.04 | P6 and P7 | <u>H-bond</u>: Asp26, His28 (**P6**) Arg88, Asn91 (**P7**)<br><u>vdW</u>: Arg88 (**P7**) |
| Eribulin | C2 | -11.87 | P6 and P7 | <u>H-bond</u>: Asp26, Asn50 (**P6**) Asn91 (**P7**)<br><u>vdW</u>: Asn50 (**P6**), Asp90 (**P7**) |
| Cydostreptin | C1 | -11.46 | P1 and P5 | <u>H-bond</u>: Tyr24 (**P1**) Lys338 (**P5**)<br><u>vdW</u>: Thr41 (**P1**) |
| Zampanolide | C2 | -11.21 | P7 | <u>H-bond</u>: Asp90, Asn91 (**P7**)<br><u>vdW</u>: His28 (**P6**) Arg88, Asp90 (**P7**) |
| Docetaxel | C2 | -9.52 | P6 and P7 | <u>H-bond</u>: Asp26, Glu27, Asn50 (**P6**) Arg88, Pro89, Asn91 (**P7**)<br><u>vdW</u>: Leu42, Gln43 (**P6**), Pro89 (**P7**) |
| Zampanolide | C1 | -7.60 | P1 | <u>H-bond</u>: Gly44, Gly45 (**P1**)<br><u>vdW</u>: Pro37, Lys40 (**P1**) |

**Figure 11:**
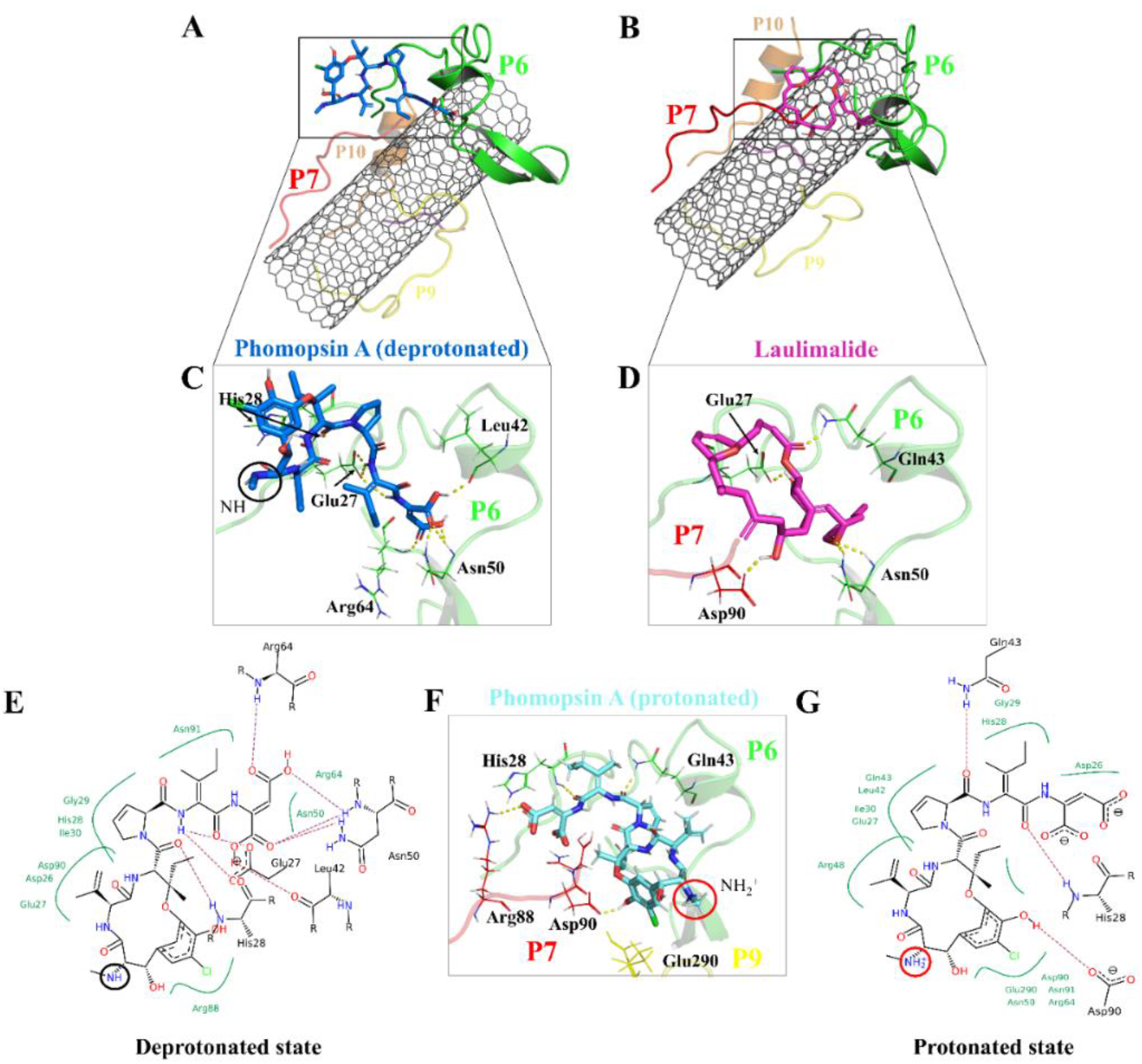
The binding profile of (**A, C**)* neutral phomopsin A (blue), (**B, D**)* laulimalide (magenta), and protonated phomopsin A (cyan). (**E, G**) Interactions of the protonated and neutral phomopsin A with the P6 and P7 amino acids. Dashed lines represent H-bonds. *: The interactions of neutral phomopsin A consisted of H-bonds to the P6: its O24 with His28 amide hydrogen; H35 with the two Glu27 carboxyl oxygen atoms; H46 with Glu27 carboxyl oxygen; H51 with Leu42 carbonyl oxygen; the O53 and O54 atoms formed weak H-bonds with Asn50 amide hydrogen; similar to O55 with Arg64. Phomopsin A formed polar interactions with Arg88 of the P7 through its O2 atom: its C28 atom affected by Gly29 backbone carbons through hydrophobic interactions similar to His28 of the P6 and the C11 atom of the ligand and Arg64 of the P6 and the C48. The protonated phomopsin A made weak H-bonders between the hydrogen of His28 (the P6) and the O45 of the ligand, similar to Gln43 (the P6) with the O34 atoms. Another hydrogen bond occurred between the NH of the guanidine in Arg88 (the P7) and the ligand’s O55; the carbonyl oxygen of Asp90 (the P7) and the H15 atom of the protonated phomopsin A. The hydrophobically interacting amino acids were Asp26 (the P6) with the C47, Glu27 (the P6) with the C36 and Leu42, and Gln43 (the P6) with the C27, where Leu 42 involved the C6 atom. Laulimalide was accommodated on the CNT by the P6 and the P7 through the H32 H-binding to the carboxyl oxygen of Asp90 (the P7), similar to the O19 atom with the amide hydrogen of Gln43 (the P6). Other involving amino acids of the P6 in the H-bond formation included Glu27 and Asn50, while His 26, Glu27, Ile49, and Gly29 bound to the protonated phomopsin A via vdW forces.

The protonated phomopsin A (at the amide nitrogen, the N1) was also docked, and the FlexX scoring function ranked it as the second-best binding energy among the studied ligands (−17.99 kJ/mol). The conformation of the protonated phomopsin A was different from its neutral structure. The ionic interaction between the protonated amine moiety (H-N1^+^) and the negatively charged oxygen atom of Glu290 in the P9 and its H-bond to the same residue caused the ligands’ conformational difference in the protonated and deprotonated states, resulting in the loss of minor interactions detected in the neutral ligand and 8.56 kJ/mol increase of binding energy (−26.56 kJ/mol vs. −17.99 kJ/mol) in the protonated form. The third strongest binding was associated with laulimalide, with −17.79 kJ/mol binding energy. (**Table 7** and **Figure 10–11**)

## 4. Conclusions

In the presented work, a microtubule-inspired non-covalent functionalization of CNTs is proposed. Ten peptides consisting of the MT’s lateral segments were studied for their potential binding to and increasing the solubility of a pristine CNT. Their effects on transforming the tube into an efficient drug carrier were also examined by analyzing the binding affinity of seventeen MT-targeting antimitotics to the functionalized CNT.

The P4 and the P9, equivalent to the M-loop of α- and β-tubulin, were bound to the CNT with the lowest LJ potential energies. The P4’s amino acid composition enabled a peptide-conformation that satisfactorily adapted the tube’s curve and resulted in a superior surface coating than the other nine peptides. However, a part of the P4 sequence was seen distant from the tube surface for ∼30 ns of the simulation time, caused by the H-bond mediating water molecules between Tyr272 and Glu284 of the P4. The intra-peptide H-bond, facilitated by Tyr272, caused steric hindrance for the remainder of the P4’s residues preventing its complete interactions to the CNT; thus, replacing Tyr272 with a Phe is expected to increase the peptide’s binding affinity. The P9 binding to the CNT, mainly half-sequence length, was regulated by its polar and charged residues. Due to the size and its specific amino acids composition, the P9 can be better accommodated by CNTs with broader surface or greater dimensions than the CNT (16,0) used in this study. In addition, replacing its Glu290 with a hydrophobic residue is suggested to eradicate the undesired polar interactions and the H-bonds with the dynamic solvent molecules. The P2 (α-H2-B3), P5 (α-H10) and P7(β-H2-B3) showed the longest retention time on the CNT, displaying more frequent interactions with the tube than the other peptides. The intra-peptide H-bonds in the P6 (β-H1-B2), such as between Tyr52 and Asn54, negatively affected the P6 binding to the CNT, costing its binding ability to the tube surface. The Asn mutation with a hydrophobic amino acid such as isoleucine can prevent the unfavorable effects. In the P1 (α-H1-B2), the equivalent segment of the P6 in β-tubulin, His28 played a critical role in strengthening the LJ potential energy due to the parallel position of His imidazole ring to the tube. However, the rotation of the Tyr’s phenyl ring under the influence of the H-bonding to Glu27, His28, and water molecules weakened the P1 binding to the CNT. A mutation at Tyr24 with Phe can omit the hydroxyl moiety’s H-bonding capability and improve the P1’s binding affinity.

The influence of the CNT’s presence on the peptides’ secondary structure was most pronounced on the P1, P6, P5, and P10. It significantly improved the P1’s configuration, stabilizing its folding as a helix and a 3-helix. Similarly, binding to the tube upgraded the configurational state of the P6 to a β-strand, more stable than the P6’. The CNT enables the P5 and P10 to maintain the same secondary structure as in microtubule, whereas their CNT-free forms, the P5’ and P10’, acted as simple coils throughout the MD trajectory.

The P2 (α-H2-B3) demonstrated a unique structural conformation by wrapping its N-terminal around the CNT in a crook-like shape. However, its Arg84 made frequent H-bonds to Arg79, Gln85, and Tyr83, pulling the P2 segment (Thr80–Thr82) away from the CNT in some MD time frames. Tyr83 and Phe87 caused spatial hindrance for the P2’s residues, preventing their complete engagement with the tube’s carbocyclic substructures. Simple hydrophobic residues, such as Leu, Val, Ile, instead of Tyr83 and Phe87, can avert the steric hindrance effect, allowing the full-length exposure of the P2 to the CNT. Mutating Arg84 with a hydrophobic amino acid is also suggested to prevent the undesired H-bonds formation. Phe87 in the P7 (β-H2-B3) blocked the peptide’s full-sequence involvements with the CNT, weakening its LJ binding energy; thus, a non-aromatic hydrophobic amino acid instead of the Phe can enhance the P7’s binding profile. The P3 (α-H4-T5) and the P8 (β-H4-T5) were the shortest peptides in sequence among the P1–P10, acted mainly as auxiliary agents to stabilize the conformational status of their adjacent peptides.

The peptides’ effects on the CNT solubility depended on the number of H-bonds each makes with the solvent molecules. Due to its residue compositions, the P6 demonstrated more hydrogen bonds with water molecules (in the C2) than the P1 (in the C1). In particular, the P2, P8, P9, and P10 showed superior potential over the P3, P4, P5, and the P7, in enhancing the CNT’s water solubility. It is noteworthy that peptides such as the P1, P4–P7, and the P9–P10, rich in polar or positively charged residues, could form strong bonds with the CNT and the solvent molecules through dynamic conformational reorientations, and thus augment CNTs solubilization. The hydrodynamic layer created by the peptides from β-tubulin in the C2 system was more widespread than the α-tubulin’s in the C1 system. In contrast, the latter formed a broader hydrophobic coating than those from the β-tubulin. Therefore, we propose a functionalization with a combination of the amphiphilic peptides from both α- and β-tubulin to improve CNTs’ aqueous solubility, cellular uptake, and efficiency to hold and transport hydrophobic drugs. That could help overcome the significant challenge in drug discovery concerning hydrophobic drugs’ insolubility, leading to inefficient bioavailability, absorption, distribution, metabolism, and excretion from the human body. The presence of the hydrophilic peptides is expected to prevent the functionalized CNT from precipitating in the blood artery allowing better fluidity in the bloodstream. Despite the tube’s frequent drift from the center of the water box up to 5.6 nm, as seen in the DCOM graph; it is considered minor, taking into account the ∼10^4^ nm diameter of the blood artery.

The drugs docking on the peptides demonstrated phomopsin A (neutral and protonated) and laulimalide as the top three best binders among the seventeen antimitotics. The two drugs were bound mainly to the P6 and the P7 of the C2 system. The charged amino acids of the P6 attracted the C-terminal charged groups of the P7 through electrostatic forces, shaping a pair-like format. In addition, the time spent on the CNT by the P6 and P7 were among the tube’s top three dwelling peptides. According to the docked ligands’ binding profiles, the P4 and the P9 (i.e., the M-loop) insignificantly participated in the ligands’ binding. In particular, the P9’s partial detachment from the CNT, in favor of H-bonds to the P6 and water molecules, indicated its imperfection as a functionalizing group. However, the long peptides in sequence, such as the P1 and the P6 with forty amino acids, or a pair of shorter peptides, such as the P6-P7, the P1-P5, and the P7-P10, can facilitate efficient conditions for a ligand binding. Remarkably, the P6-P7 pair was the most preferred binding site for several ligands, namely, phomopsin A (neutral and protonated), laulimalide, epothilone A, epothilone D, discodermolide, eribulin, and docetaxel.

The designed CNT-peptides complex is deemed a more biocompatible drug carrier and is more likely to increase CNT’s cellular uptake or modulate neutrophil activation in the bloodstream than its pristine form, considering the improved binding capacity of the peptide-coated tube to the plasma proteins surface. The proposed functionalization approach is expected to help advance *in vitro* and *in vivo* attempts to design and develop bioapplicable and biocompatible functionalized CNTs.

## Acknowledgments

The authors acknowledge funding support from the Natural Sciences and Engineering Research Council of Canada (NSERC), Discovery Grant (No. 212654), awarded to L. A. The authors thank Compute Canada for accessing the HPC facilities and the support from the technical staff of ACENET.

